# TRIM9 controls growth cone responses to netrin through DCC and UNC5C

**DOI:** 10.1101/2024.05.08.593135

**Authors:** Sampada P. Mutalik, Ellen C. O’Shaughnessy, Chris T. Ho, Stephanie L. Gupton

## Abstract

The guidance cue netrin-1 promotes both growth cone attraction and growth cone repulsion. How netrin-1 elicits these diverse axonal responses, beyond engaging the attractive receptor DCC and repulsive receptors of the UNC5 family, remains elusive. Here we demonstrate that murine netrin-1 induces biphasic axonal responses in cortical neurons: attraction at lower concentrations and repulsion at higher concentrations using both a microfluidic-based netrin-1 gradient and bath application of netrin-1. TRIM9 is a brain-enriched E3 ubiquitin ligase previously shown to bind and cluster the attractive receptor DCC at the plasma membrane and regulate netrin-dependent attractive responses. However, whether TRIM9 also regulated repulsive responses to netrin-1 remained to be seen. In this study, we show that TRIM9 localizes and interacts with both the attractive netrin receptor DCC and the repulsive netrin receptor, UNC5C, and that deletion of murine Trim9 alters both attractive and repulsive responses to murine netrin-1. TRIM9 was required for netrin-1-dependent changes in surface levels of DCC and total levels of UNC5C in the growth cone during morphogenesis. We demonstrate that DCC at the membrane regulates growth cone area and show that TRIM9 negatively regulates FAK activity in the absence of netrin-1. We investigate membrane dynamics of the UNC5C receptor using pH-mScarlet fused to the extracellular domain of UNC5C. Minutes after netrin addition, levels of UNC5C at the plasma membrane drop in a TRIM9-independent fashion, however TRIM9 regulated the mobility of UNC5C in the plasma membrane in the absence of netrin-1. Together this work demonstrates that TRIM9 interacts with and regulates both DCC and UNC5C during attractive and repulsive axonal responses to netrin-1.

## Introduction

Netrin-dependent axon guidance is crucial for early neuronal development; deletion of the gene that encodes netrin-1 (*Ntn1*) causes embryonic lethality (Kennedy et al., 1994; Bin et al., 2015). *Ntn1* deletion is associated with several axon guidance defects including agenesis of the corpus callosum, absence of hippocampal and spinal commissures, and disorganized thalamocortical projections (Mitchell et al., 1996; Braisted et al., 2000; Bradford et al., 2009; Bin et al., 2015). Interestingly, netrin-1 induces both attractive and repulsive responses *in vivo* (Keino-Masu et al., 1996; Srivatsa et al., 2014; Dailey-Krempel et al., 2023), but the mechanisms mediating these diverse responses are not fully understood (Boyer and Gupton, 2018). Modeling results *in vivo*, we previously found that a gradient of chicken netrin-1 induced biphasic axon turning responses *in vitro*, which was dependent upon the local concentration of netrin-1, with attractive axon turning occurring at lower concentrations of a netrin gradient and repulsive axon turning occurring at higher concentrations (Taylor et al., 2015; Boyer et al., 2019).

Engagement with distinct receptors or combinations of receptors is one potential mechanism generating diverse axonal responses to netrin. Multiple netrin receptors including DCC (Keino-Masu et al., 1996), UNC5 family members (Leonardo et al., 1997), and DSCAM (Liu et al., 2009) regulate netrin-dependent growth cone motility, attractive and repulsive axon turning, and axon branching (Moore et al., 2007; Rajasekharan and Kennedy, 2009; Boyer and Gupton, 2018). Like deletion of *Ntn1*, deletion of *Dcc* also results in agenesis of corpus callosum (Fothergill et al., 2014). Whereas multiple studies have shown functions of different UNC5 family members (A-D) in cerebellar development (Ackerman et al., 1997; Goldowitz et al., 2000; Purohit et al., 2012). UNC5C is also expressed in the developing cortex and its deletion is postnatally lethal (Kim and Ackerman, 2011). Interestingly, UNC5C and DCC regulate netrin-dependent corpus callosum development and thalamocortical projections by promoting both repulsive and attractive axon responses (Srivatsa et al., 2014; Powell et al., 2008), but the cellular and molecular mechanisms downstream of netrin receptors that distinguish attractive and repulsive responses are unknown.

Netrin receptors engage downstream signaling pathways to regulate cytoskeletal dynamics and membrane remodeling during morphogenesis. We have previously shown that DCC is ubiquitylated in the absence of netrin-1, and that this modification is reversed in the presence of netrin-1. Focal adhesion kinase (FAK) is a key kinase activated in response to netrin-1. In netrin-dependent attractive responses, FAK undergoes autophosphorylation at tyrosine 397 (pY397), phosphorylates the cytoplasmic tail of DCC, and recruits Src kinase and downstream cytoskeletal regulators (Li et al., 2004; Ren et al., 2004; Moore et al., 2012; Plooster et al., 2017; Mahadik and Lundquist, 2023). Presumably downstream signaling during attractive and repulsive responses are distinct, but in the case of FAK, the nature of this dichotomy is not known. We previously identified the brain-enriched E3 ubiquitin ligase TRIM9 as an interaction partner of DCC that regulates netrin-dependent attractive axon turning, axon branching, and synaptogenesis in murine neurons (Winkle et al., 2014; Menon et al., 2015; Plooster et al., 2017; Boyer et al., 2019; Menon et al., 2021; McCormick et al., 2024). We found that in the absence of netrin-1, TRIM9 was required for ubiquitylation of DCC, which blocked FAK-mediated phosphorylation of DCC, and that in the absence of Trim9, DCC was hyperphosphorylated and responses to netrin-1 were ablated (Plooster et al., 2017). We also found that deletion of murine Trim9 resulted in increased thickness of corpus callosum (Winkle et al., 2014). This phenotype was associated with increased aberrant axon branching in the callosum (Winkle et al., 2014), along with increased projection to the internal capsule (Menon et al., 2015). Interestingly, *Unc5c* deletion showed similar defects in the internal capsule *in vivo* (Srivatsa et al., 2014), suggesting that TRIM9 and UNC5C may collaborate during netrin-dependent repulsive responses.

In the current study, we show that murine netrin-1 induces a concentration-dependent biphasic growth cone response, with attractive turning and growth cone expansion occurring at lower concentrations of netrin-1, and repulsive turning and growth cone retraction and collapse occurring at higher concentrations of netrin-1. Further, we identified TRIM9 as a regulator of both attractive and repulsive netrin-dependent biphasic responses. Our findings suggest that TRIM9 regulates distinct guidance responses through the differential regulation of netrin receptors DCC and UNC5C at attractive and repulsive concentrations of netrin-1. We show that TRIM9 interacts with both UNC5C and DCC. We show that TRIM9 regulates DCC and UNC5 levels in growth cones, and downstream netrin-dependent changes in FAK activation. Using immunocytochemistry and live imaging approaches, we show that lack of netrin responsiveness in *Trim9* ^-/-^ neurons stems at least partially from deregulation of surface receptors DCC and UNC5C.

## Materials and Methods

### Animals

Mouse lines were on a C57BL/6J background and bred at the University of North Carolina at Chapel Hill (UNC) with the approval from the Institutional Animal Care and Use Committee. Generation of the *Trim9* ^-/-^ mice was described previously (Winkle et al., 2014). Timed pregnancies were set by placing male and female mice together overnight. Next day is called E 0.5 if the female had a vaginal plug.

### Plasmids, antibodies and reagents

Doxycycline (Sigma, D9891)-inducible adenoviral constructs were generated for Halo-TRIM9 and UNC5C-pHmScarlet using the adeno-X™ system 3 (Takara Bio, 631180), using the detailed protocol outlined by (O’Shaughnessy et al., 2024). The TRIM9-Halo construct was created using pcs2-Myc-TRIM9 plasmid (Winkle et al., 2014). The UNC5C-pHmScarlet was created using pCDNA3-UNC5CHA (Guofa Liu, University of Toledo) and pH-sensitive mScarlet0 (Addgene, 166891), positioning mScarlet0 after the Ig2 domain within the extracellular region of UNC5C. For colocalization experiments, the previously described TRIM9△RING-mCherry mutant, lacking the ligase domain, was utilized (Winkle et al., 2014). Janelia Fluor 646 HaloTag ligand (Promega, GA1120) was used for labeling of Halo-tagged proteins. Immunoprecipitation experiments utilized pEGFP-N1 (Clontech, 6085-1), DCC-pHluorin (Plooster et al., 2017), pcs2-kennedy, pcs2-myc, and pcs2-DCC-HA (Winkle et al., 2014). The Myc-tagged mouse netrin-1 construct was obtained from Samantha Butler’s lab (UCLA). Recombinant mouse myc-netrin-1 was cloned into pSBI-GN vector (Addgene, 60517) to prepare a stable HEK293 cell line expressing mouse netrin-1.

Myc monoclonal antibody 9E10 (UNC antibody core), antipFAKpY397 (44-625G,ThermoScientific), anti-total FAK (T44-625G, ThermoScientific), rabbit polyclonal anti-GFP (A11122, ThermoScientific), Anti-HA (05-904, Millipore), 680LT goat anti-rabbit antibody (926-68021, Licor), 800CW donkey anti-mouse antibody (925-32212, Licor) were used for immunoblotting. DCC goat polyclonal antibody (PA5-47951, ThermoFisher Scientific), UNC5C rabbit polyclonal antibody (PA5-95332, ThermoFisher Scientific), mouse *β*-III tubulin antibody (801202, Bioligand), Alexa568 phalloidin (A12380, ThermoScientific), Alexa488 anti-goat (A32814TR, Thermo-Scientific), Alexa647 anti-rabbit (A32795, ThermoScientific), and Alexa405 anti-mouse (A48257, ThermoScientific) were used in immunocytochemistry experiments. In addition, anti-goat 680 (Biotium, 20196-1) was used for staining DCC in STED experiments, DCC goat polyclonal antibody (PA5-47951, ThermoFisher Scientific) was also used for function blocking experiments.

### Netrin preparation

Mouse netrin-1 was concentrated from conditioned media of stably expressing HEK293 cell lines. Briefly, HEK293 cells stably expressing myc-netrin-1 were cultured in DMEM media supplemented with 10% FBS and Neomycin (20 ug/ml, N1142, Sigma) to 80-90% confluency. Cells were washed with sterile, warm PBS followed by incubation in Opti-MEM (31985070, Gibco) overnight. Secreted netrin-1 in Opti-MEM was concentrated using Amicon filters tubes with a 30 kDa cut-off (UFC903008, Millipore). The concentration of netrin-1 was estimated by SDS-PAGE using Coomassie staining with BSA as a standard.

### Neuronal culture and imaging experiments

Cortices from E15.5 litters of *Trim9* ^+/+^ and *Trim9* ^-/-^ pups were dissected and dissociated with 1X trypsin (20 min at 37°C) (Winkle et al., 2014). Neurons were plated on PDL coated surfaces in neurobasal medium supplemented with B27 and Glutamax. In a 35 mm dish, 600,000 neurons were plated on 25 mm coverslip for growth cone response imaging experiments.

### Immunocytochemical growth cone analysis

At 2 days in vitro (DIV), neuronal cultures were treated with the indicated concentration of murine netrin-1 or media sham. After one hr, neurons were fixed in 37°C PHEM buffer (60 mM PIPES (pH: 7), 25 mM HEPES (pH: 7), 10mM EGTA (pH: 8), 2 mM MgCl_2_, 0.12 M sucrose) with 4% paraformaldehyde (PFA) for 20 min. DCC goat polyclonal antibody (PA5-47951, ThermoFisher Scientific, 1:1000) was added after 2 DIV, followed by fixation next day. Membrane DCC was labeled using a DCC goat polyclonal antibody (1:200) for 1 hr before permeabilization, followed by three 5 min washes with PBS. Cells were stained with an anti-goat Alexa 488 antibody, followed by additional PBS washes. Permeabilization was achieved with 0.2% Triton X in PBS for 5 mins, followed by PBS washes. After blocking with 10% normal donkey serum (NDS) for 1 hr, cells were incubated with anti UNC5C (1:200) and *β*-III tubulin (1:1000) antibodies for 1 hr, three 5 min washes, followed by incubation with secondary antibodies (anti-mouse Alexa 405, anti-rabbit Alexa 647, and phalloidin Alexa568. Coverslips were mounted using mounting media (20 mM Tris pH: 8, 0.5% N-propyl gallate, and 90% glycerol) and sealed with nailpolish. Growth cones were imaged on an Olympus IX83-ZDC2 using 100X, 1.50 NA TIRF objective and ORCA FUSION BT sCMOS. Image analysis was performed in FIJI (ImageJ). Images were blinded using blind analysis tools in FIJI. Growth cone area was measured using polygon selection tool and compared using Mann-Whitney test in GraphPad Prism9.

### Transduction and STED imaging

At 1 DIV, *Trim9* ^-/-^ cortical neurons (600,000 cells) plated on 25 mm coverslip were transduced with adenoviral TRIM9 -Halo (5 ul). Doxycycline (final concentration: 1 ug/ml) was added at the time of transduction to induce expression. At 3 DIV, neurons were incubated with 200 nM Janelia Fluor 646 HaloTag ligand for 15 mins, followed by three washes with the culture media. Cultures were fixed and stained for DCC at the cell surface and total UNC5C as described above, except with donkey anti-goat 680 and donkey anti-rabbit. ProLongX™ Diamond Antifade Mountant (P36965, ThermoScientific) was used for mounting coverslips per the manufacturer’s protocol. STED imaging was carried out using a Leica STELLARIS 8 FALCON STED nanoscope equipped with APO CS2 100x, NA:1.4 oil objective. For TRIM9-Halo, endogenous DCC and UNC5C: 590 nm (intensity: 5%), 637 nm (intensity: 20%) and 685 nm (intensity: 20%) excitation lasers were used, respectively. Three color STED was performed using 775 nm (intensity: 90%) pulse depletion laser. Images were acquired by line scanning with 33 nm resolution, each line was scanned four times.

### Time lapse imaging and analysis of growth cone dynamics

*Trim9* ^+/+^ and *Trim9* ^-/-^ cortical neurons (600,000 cells) were plated in 35 mm Cellivis (D35-20-1.5-N) with 20 mm microwell movie dishes and imaged over time in Differential Image Contrast (DIC) mode before and after netrin-1 (1200 ng/ml) addition for 1 hr on Olympus IX83-ZDC2 equipped with Tokai-hit incubator, maintained at 37°C with 5% CO2 and ORCA FUSION BT sCMOS camera with a 100X, 1.51 NA objective. Images were acquired using CellSens every 10 min using multi-acquisition tool. Time lapse images were blinded, and growth cones were classified into three categories: ruffling or advancing growth cones (+), paused growth cones (=), and collapsed or retracting growth cones (-). The percentage of growth cone states were compared in GraphPad Prism9 using Fisher’s *χ* test.

### Axon turning

Microfluidic devices were prepared as per the published protocol (Taylor et al., 2015). Prepolymer and catalyst (Sylguard, DC4019862) were mixed at 10:1 ratio; using a plastic knife. 10g PDMS was added per device, degassed in desiccator for 2-3 hr, and then cured at 65°C at least overnight before the use. The day before culturing, devices were cut and assembled on PDL-coated coverslips and incubated with neuronal culture media. 10 μl of 18 million neurons/ml were seeded in cell compartment. Once axons had extended through the microchannels into the axon viewing chamber, 48 to 72 hrs, a gradient was established. To do this, murine netrin-1 (1200 ng/ml) + rhodamine dextran (0.5 ug/ml, 10,000 kDa, sigma R9379) or only rhodamine dextran gradient was added to source chamber one, and 50-75 μl of media was removed from the sink chamber, as previously described (Taylor et al., 2015). Images of the gradient and axons in the axon imaging chamber were acquired on an ORCA FUSION BT sCMOS camera with a 20X, NA: 0.85 objective by widefield fluorescence and DIC, respectively on an Olympus IX83-ZDC2 equipped with Tokai-hit incubator maintained at 37°C with 5% CO2. Tiled images of the entire axon chamber were acquired every 5 mins, a tiled image of the fluorescent dextran gradient was acquired every hr. Individual devices were images up to 24 hrs, to obtain maximum number of axonal entry and turning behaviors in axon viewing area. High and low netrin regions were determined from the dye field images for each device. The gradient was stable over the course of imaging and thus a mean projection over time was used to reduce noise. An ROI across the entire field (in the x-direction) was selected to remove any debris or cells which distorts fluorescence. The mean intensity in the x-dimension of the ROI was plotted as a function of x-position in the device. The boundary between high and low netrin was set just above where the dye intensity plateaued to the low baseline. Axons were classified into “high” and “low” netrin based on the initial x-position of the growth cone.

Images were analyzed with a custom script in ImageTank (O’Shaughnessy et al., 2019), (https://www.visualdatatools.com/ImageTank/) to quantify growth parameters for all axons meeting predefined criteria: axons were not considered for the analysis if they grew along other axons, crossed other axons, interacted with the edges of the viewing window, or did not remain healthy for at least 2 hrs (tracking was terminated at 10 hrs). Branches were included in our analysis if they clearly separated from the primary axon and met the criteria outlined for inclusion above. After the starting and ending time points were determined, the images were manually corrected for drift or shifts in image stitching if needed. For both the starting and ending frames the angle of growth was measured relative to a horizontal line drawn down the gradient. The first segment of the angle was the horizontal line ending on the stalk of the axon and the second segment connected the axon stalk to the middle of the growth cone. Using these six points (three from the starting angle and three from the ending angle) we measured: distance traveled (shortest distance between growth cone at the start and stop), velocity, initial and final angles of growth, difference in growth angle, the initial and final directions with respect to the gradient and axon retraction. When comparing the difference in growth angle we excluded axons that retracted or that failed to grow more than 5 μm over the duration of imaging as these axons have stalled and the angle is not meaningful. For axons that retracted we considered the velocity to be negative and these and all stalled axons were included in the velocity measurements. Compiled data were compared in Graphpad Prism 9 using unpaired t test.

### FRAP experiments and analysis

Cortical neurons (600000 cells) from *Trim9* ^+/+^ and *Trim9* ^-/-^ embryos were cultured in35 mm Cellivis (D35-20-1.5-N) with 20 mm microwell. At 2 DIV, neurons were transduced with 5 ul UNC5C-mscarlet adenovirus and media supplement with doxycycline, final concentration: 1 ug/ml for the induction. For FRAP experiments, neurons were imaged on an Olympus IX83-ZDC2 equipped with Tokai-hit incubator, maintained at 37°C with 5% CO2 using a 100X, 1.51 NA objective, in TIRF mode (penetration depth: 110 nm) using ORCA FUSION BT sCMOS camera. Initially, images were captured every 250 ms followed by bleaching in wide field using 405 laser using 100% laser power. Post-bleaching, recovery of fluorescence was captured every 250 ms. 1200 ng/ml netrin-1 was added, and post netrin FRAP experiment was performed a different region in the same cell within 30 mins of netrin-1 addition. FRAP data were analyzed as described (Day et al., 2012). Images were background subtracted and fluorescence intensity of the whole axon neurite was measured. A region of interest was drawn around bleached circular region in 405 channel and fluorescence was recorded across the time lapse using multi-measure tool in FIJI. Normalized FRAP data was calculated as follows:

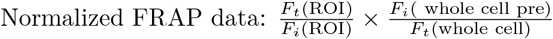

In this method, loss of molecules due to the photo-bleaching is corrected by simple ratio method, in which bleaching ROI intensity is divided by whole cell intensity in the equation above. Recovery curves were generated and fitted to single exponential curves in GraphPAd Prism 9. If the R^2^ was *>*0.6, data were considered in class averages. The recovery rates and half time across treatments were compared in GraphPad Prism 9. Wilcoxon test for paired comparisons before and after netrin-1. Half times of recovery in *Trim9* ^+/+^ and *Trim9* ^-/-^ were compared using Mann Whitney test.

### Membrane dynamics of UNC5C

Cortical neurons (600,00 cells) from *Trim9* ^+/+^ and *Trim9* ^-/-^ embryos were cultured in35 mm Cellivis (D35-20-1.5-N) with 20 mm microwell. At 2 DIV, neurons were transduced with 5 μl UNC5C-pH-mScarlet adenovirus and media was supplemented with doxycycline, final concentration: 1ug/ml for induction. At 48 hrs post-transduction, images were acquired before netrin application followed by every 10, 20, 40 and 60 mins post-netrin (1200 ng/ml) application on an Olympus IX83-ZDC2 equipped with Tokai-hit incubator, maintained at 37°C with 5% CO2 using a 100X, 1.51 NA objective, in TIRF mode (penetration depth: 110 nm) using ORCA FUSION BT sCMOS camera. Images were imported, background subtracted and thresholded using auto thresholding in Image J. Region of interests were saved in ROI manager. Mean intensities were measured and fold change in mean intensity after netrin addition was compared using Kruskal Wallis test with Dunn’s correction, plotted in GraphPad Prism9.

### Coimmunoprecipitation and immunoblotting

For DCC-UNC5C interaction studies, previously developed HEK293 *TRIM9* ^-/-^ cell line was used (Menon et al., 2015). *TRIM9* ^+/+^ and *TRIM9* ^-/-^ HEK293 cells were cultured in DMEM + 10% FBS at 37°C overnight followed by transfection with plasmids encoding DCC-HA and UNC5C-GFP using lipofectamine 2000 per manufacture protocol. After overnight expression, cells were treated with murine netrin-1 (1200 ng/ml) or media sham for one hr. *TRIM9* ^+/+^ and *TRIM9* ^-/-^ HEK293 cells expressing pEGFP-N1 with DCC-HA were used as a control. Cells were lysed in lysis buffer (50 mM Tris, pH 7.5, 200 mM NaCl, 2 mM MgCl_2_, 10% glycerol, 1% NP-40, supplemented with 15 mM sodium pyrophosphate, 50 mM NaF, 40 mM *β*-glycophosphate, 1 uM PMSF, 1 mM sodium vanadate. 1 ug/ml leupeptin, 5 ug/ml aprotinin). UNC5C-GFP was precipitated using GFP trap beads (ChromoTek, gta) for 1 hr at 4°C with agitation. Beads were washed three times with lysis buffer and boiled with sample buffer. Samples were resolved by SDS-PAGE. DCC-HA and UNC5C GFP in the immunoprecipitates were analyzed by immunoblotting using anti-HA and anti-GFP antibodies. Anti-mouse 800 and anti rabbit 680 secondary antibodies were used for immunolabeling.

To evaluate whether TRIM9 interacts with DCC and UNC5C, *TRIM9* ^-/-^ HEK 293 cells were cultured in DMEM+10% FBS at 37°C overnight followed by transfection with lipofectamine 2000 with plasmids. Myc-TRIM9△RING was expressed with DCC-pHluorin and UNC5C-GFP in *TRIM9* ^-/-^ HEK293 cells. *TRIM9* ^-/-^ HEK293 cells expressing Myc, DCC-pHluorin and UNC5C-GFP were used as a control. Myc-TRIM9△RING was precipitated using myc antibody for overnight at 4°C on rocker, followed by isolation using Protein-A-Agarose beads (sc-2001, Santa Cruz) for three hrs at 4°C on the rocker. Beads were washed three times with lysis buffer and boiled in sample buffer, followed by SDS-PAGE and immunoblotting with anti-myc and anti-GFP primary antibodies. Anti-mouse 800 and anti rabbit 680 secondary antibodies were used for immuno-labeling.

For pFAK analysis (pY397), cortical neurons were cultured (3 million neurons/9.6 cm^2^ well) from Trim9^+/+^ and Trim9^-/-^ mice for 2 DIV. Sham treatment, 600 ng/ml netrin-1, or 1200 ng/ml netrin-1 was bath applied for 10 mins. Neurons were lysed using Radio Immunoprecipitation Assay (RIPA) buffer (150 mM NaCl, 1% NP-40, 0.5% sodium deoxycholate, 0.1% SDS, 50 mM Tris, pH: 8, supplemented with 15 mM sodium pyrophosphate, 50 mM NaF, 40 mM *β*-glycerophosphate, 1 μM PMSF, 1 mM sodium vanadate, 1 μg/ml leupeptin, and 5 μg/ml aprotinin). Lysates were boiled in sample buffer, followed by SDS-PAGE and immunoblotting using pFAK and total FAK antibodies and GAPDH antibodies for a loading control. Anti-mouse 800 and anti-rabbit 680 secondary antibodies were used for detection.

All the Blots were imaged in Odyssey Imager (Licor). Immunoblots were analyzed using Image studio (version: 5.2.5). Blots were background subtracted followed by measuring intensity by drawing rectangles around the bands. Co-immunoprecipitation ratios were normalized to the sum of the measurements within the blots. Each ratio was divided by sum total of the ratio (Degasperi et al., 2014). pFAK/total FAK ratio were normalized similarly. Data were plotted and compared using Mann-Whitney test in GraphPad Prism 9.

## Results

### TRIM9 is required for netrin-dependent attractive and repulsive axon turning

The growth cone at the tip of the extending axon is critical for sensing guidance cues, such as netrin-1. For historical purposes, the axon guidance field has utilized chicken netrin-1 to study axonal responses to netrin (Kennedy et al., 1994; Serafini et al., 1996). We previously found that axon turning in a netrin-1 gradient was biphasic, with attractive turning occurring at low concentrations of chicken netrin-1, and repulsive turning occurring at high concentrations (Menon et al., 2015; Taylor et al., 2015). Here we wanted to examine if murine netrin-1 exhibited similar regulation of axonal behavior. We first used growth cone area after acute bath application of increasing concentrations of murine netrin-1 (1 hr) to evaluate concentration-dependent responses (Figure S1A-B). Previous work found that a concentration of 600 ng/ml of chick netrin-1 induced growth cone expansion (Menon et al., 2015; Boyer et al., 2019). We found similarly that growth cone area increased after addition of 600 ng/ml murine netrin-1, whereas at concentrations of 1200 ng/ml and above, growth cone area decreased, suggestive of growth cone collapse. This suggested murine netrin-1 also promoted concentration dependent biphasic responses.

To better define if these were attractive and repulsive responses to netrin-1, we wanted to determine if a gradient of murine netrin-1 induced axon turning. We previously developed a microfluidic device to assess axon turning of slow growing murine axons over many hrs ((Figure 1A). In this device axons grow into a fluidically isolated chamber where we rapidly induce a stable, reproducible gradient of an axon guidance cue and fluorescent dextran through a simple pipetting procedure (Taylor et al., 2015). Previously we found that axons of embryonic cortical neurons exhibited a biphasic response in a gradient of chicken netrin-1. At the low end of the gradient, attractive turning was observed, whereas at the high end of the gradient, repulsive turning was observed (Menon et al., 2015; Taylor et al., 2015). We examined axon turning behavior of cortical neurons in a gradient of rhodamine dextran and murine netrin-1 (source concentration 1200 ng/ml, (Figure 1B, Videos 1-2). We observed significantly different axon turning (p*<*.001) and growth velocity (p = .02) at the low and high ends of a murine netrin-1 gradient (Figure 1C, D). At the low end of the gradient, axons grew faster and turned towards the netrin (positive turning angles). In contrast at the higher end of the gradient, axons grew slower and turned away from the netrin source (negative turning angles). Examination of the percentage of axons growing toward (62.5%) or away from netrin (85.2%) at the low and high end of the gradient, respectively further supports this biphasic response (Figure S1C). Further analysis of final growth direction and axon turning angles suggests that turning angle may underestimate axonal response to a netrin-1 gradient, in cases where axons were randomly oriented prior to establishing a netrin-1 gradient and showed small turning angles (Figure S1C). These turning and growth responses were netrin-dependent, as a gradient of dextran alone did not induce biphasic turning (p=.92), nor influence growth velocities (p=.21, Figure 1C,D). In addition, most axons in a dextran gradient showed very small turning responses (FigureS1C). Thus, consistent with chicken netrin-1, mouse netrin-1 induced concentration-dependent biphasic responses.

**Figure 1:**
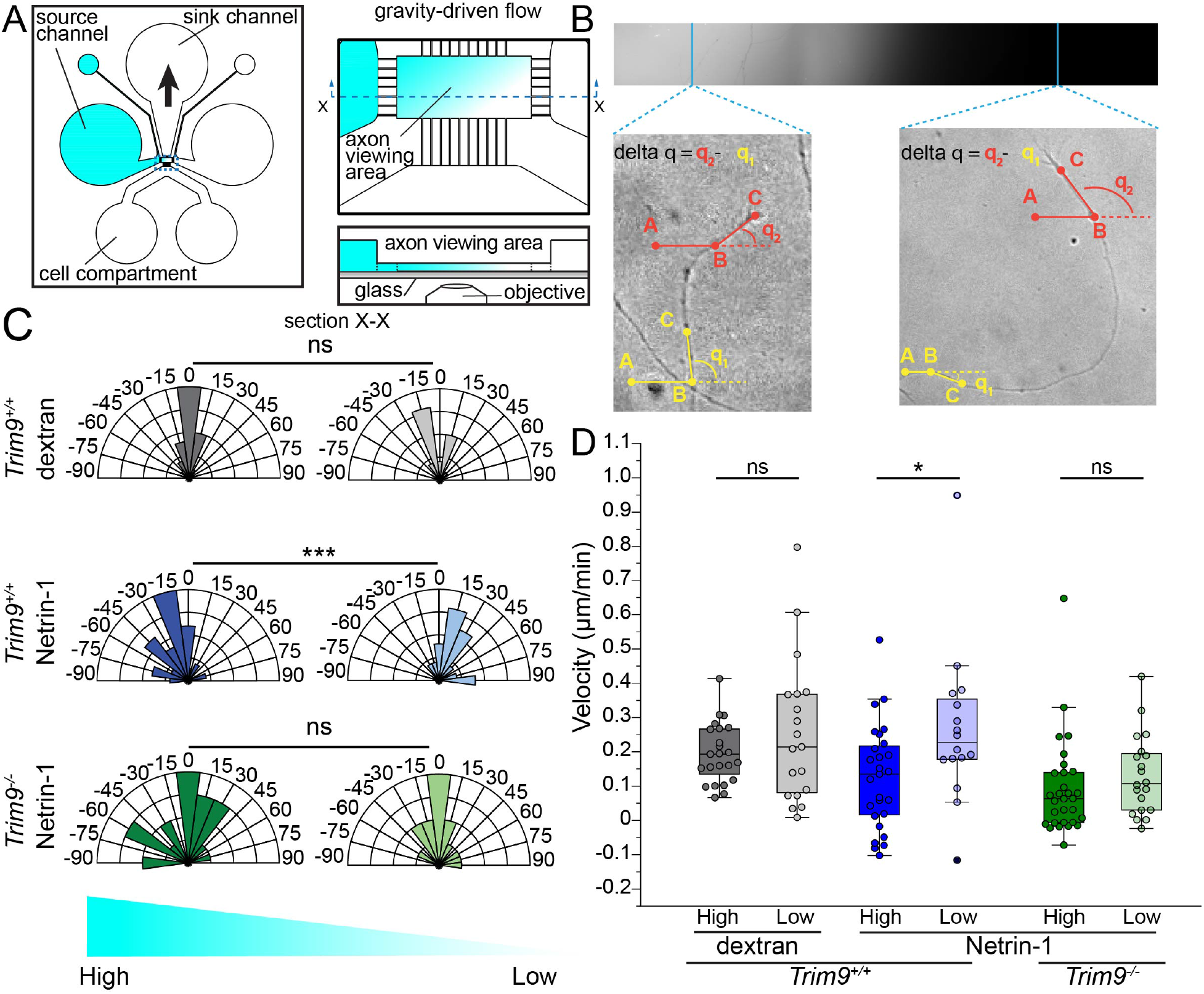
Netrin-dependent biphasic turning responses. **A**. Schematic of microfluidic device. The left source channel is filled with 150 μl of neuronal growth media supplemented with dextran + murine netrin-1 (1200 ng/ml), other wells are filled with neuronal growth media. 50-75 ul of media is removed from the sink channel to generate gravity driven flow and a stable gradient is established in the axon viewing area. **B**. Example of gradient generated across the axon viewing chamber and *Trim9* ^+/+^ cortical axons turning away and toward the high and low end of the netrin gradient, respectively. Analysis procedure for defining axon turning angle is demonstrated. **C**. Axon turning angles (°) measured at high and low end of dextran and netrin-1 gradients in indicated genotypes. Angles *>* 90° or *<*-90° are grouped into 90° and -90°, respectively. p values are from two-tailed, unpaired t test. *** indicates p *<*.001, ns indicate p *>* 0.05. **D**. Velocities of growth cone extension of *Trim9* ^+/+^ and*Trim9* ^-/-^ cortical axons in response to dextran or netrin-1 + dextran gradient. More than 10 axons per condition. p values are from two-tailed, unpaired t test. * indicates p *<* .05, ns indicate p *>* 0.05. Please refer to videos 1-2.

In our earlier studies we found that the brain-enriched E3 ubiquitin ligase TRIM9 was required for attractive axon turning (Menon et al., 2015), but we did not report a role for TRIM9 in repulsive axon turning. In the microfluidic device, axonal turning behavior (p=.26) and growth rate (p=.3) of *Trim9* ^-/-^ cortical axons was not significantly different in the high and low end of a netrin-1 gradient (Figure 1C,D). This indicated TRIM9 was required for netrin-dependent biphasic turning behavior and changes in growth speed.

### TRIM9 is required for attractive and repulsive growth cone responses to netrin-1

For subsequent experiments, we bath applied 600 ng/ml (low) or 1200 ng/ml (high) murine netrin-1 to model netrin-dependent attractive and repulsive responses, respectively. To examine if TRIM9 was required for attractive and repulsive responses, we performed experiments in *Trim9* ^+/+^ and *Trim9* ^-/-^ neurons (Figure 2A). Consistent with the previous results (Menon et al., 2015), the baseline growth cone area from *Trim9* ^-/-^ neurons was larger compared to axonal growth cones of *Trim9* ^+/+^ neurons (Figure 2B, *p <* 0.05). *Trim9* ^+/+^ neurons exhibited a biphasic growth cone response to netrin: low netrin-1 treatment increased growth cone area (Figure 2B, *p <* 0.0001)., whereas high netrin-1 decreased growth cone area (Figure 2B, p *<* 0.05), compared to media sham control. This netrin-dependent biphasic growth cone response was absent in *Trim9* ^-/-^ neurons (Figure 2B). This indicated that both attractive and repulsive growth cone responses to murine netrin-1 required TRIM9. Collectively, acute netrin treatment and netrin-gradient experiments suggested that TRIM9 regulated concentration-dependent attractive and repulsive axonal responses to murine netrin-1.

**Figure 2:**
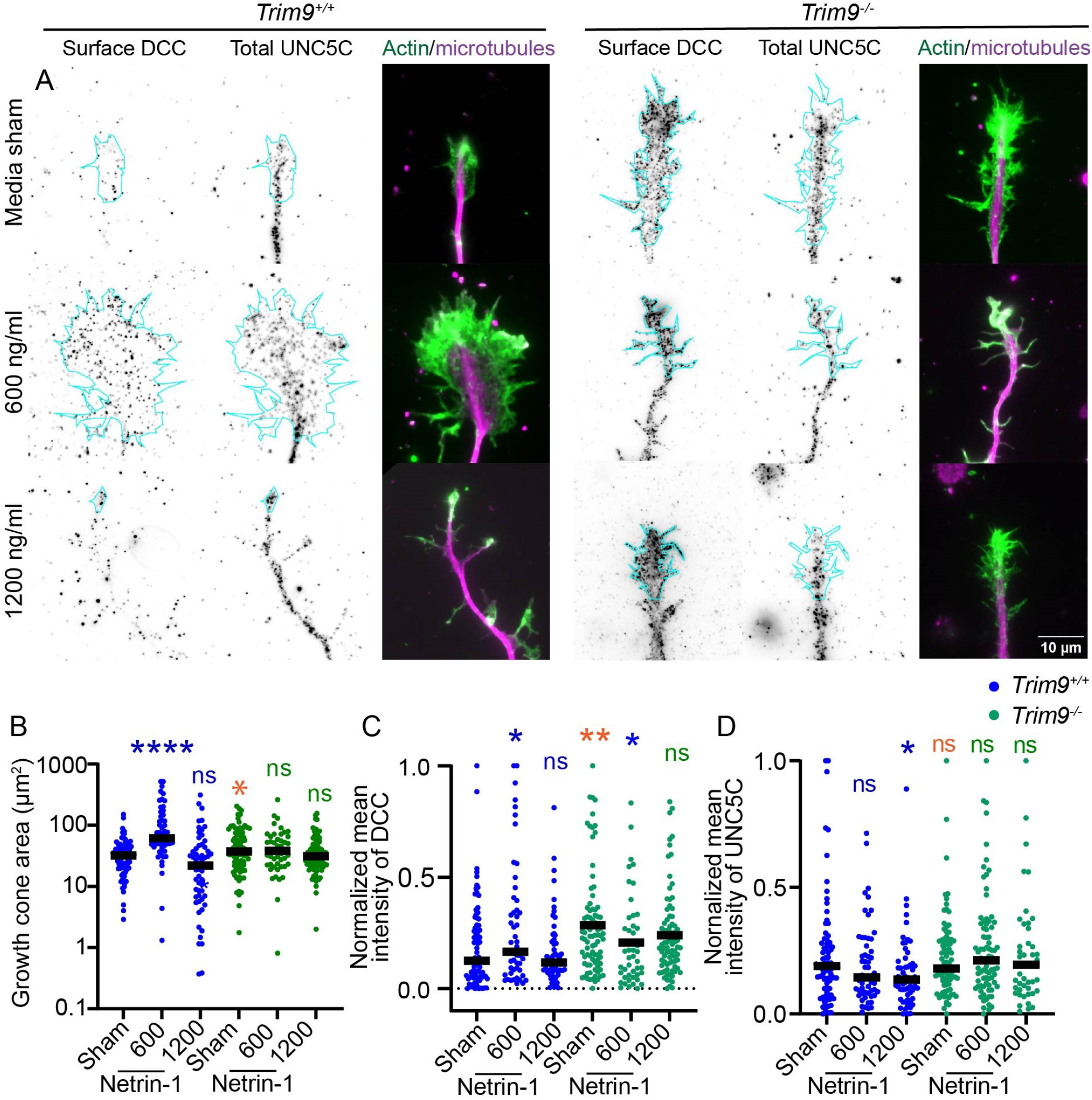
TRIM9 is required for concentration-dependent attractive and repulsive growth cone responses to netrin-1. **A**. Immunostaining of surface levels of DCC and total levels of UNC5C in *Trim9* ^+/+^ and *Trim9* ^-/-^ growth cones after application of media sham, 600 ng/ml, and 1200 ng/ml of netrin-1. F-actin is stained with phalloidin (green) and microtubules (magenta) with *β*-III tubulin antibody. **B**. Quantification of growth cone area. **C**. Quantification of membrane levels of DCC in the growth cone. **D**. Quantification of total UNC5C levels in the growth cone. The number of experiments is four biological replicates, with *>* 50 growth cones per condition. The p-values are from the Mann-Whitney test. Statistical comparisons between *Trim9* ^+/+^ and *Trim9* ^-/-^ growth cones are shown in orange, comparisons of treatment condition in *Trim9* ^+/+^ growth cones are shown in blue, and comparisons of treatment condition in *Trim9* ^-/-^ growth cones are shown in green.

To independently evaluate netrin-induced repulsion at the high netrin-1 concentration, we performed time-lapse imaging of growth cones before and after acute bath application of 1200 ng/ml netrin-1. We classified growth cones into three categories: ruffling or advancing growth cones (+), paused growth cones (=), and collapsed growth cones or retracting axons (-) (Figure S2, Video 3 and 4). Netrin-1 application significantly changed the distribution of growth cones in these categories in both *Trim9* ^+/+^ neurons (p *<* .0001) and *Trim9* ^-/-^ neurons (p *<* .004). The percentage of collapsed/retracting growth cones increased from 11.2% to 42.8% upon netrin-1 application in *Trim9* ^+/+^ neurons (Figure S2, Video 3, 4). The percentage of collapsed/retracting growth cones only increased from 4.9% to 11.34% in *Trim9* ^-/-^ neurons.

DCC and UNC5 are appreciated to function as attractive and repulsive receptors for netrin-1 (Boyer and Gupton, 2018). Our previous studies demonstrated that murine DCC clustered at the plasma membrane in response to low concentrations of chicken netrin-1 (Plooster et al., 2017). The interpretation of this was slightly complicated, as there is no homolog for DCC in the chicken genome, only the paralog neogenin (Phan et al., 2011). As such, we wondered how DCC levels at the neuronal surface were altered by different concentrations of murine netrin-1. To assess levels of DCC at the plasma membrane, we stained cortical neurons prior to permeabilization with an antibody that recognizes an extracellular epitope on DCC (Figure 2A) after 1 hr of sham, low, or high netrin-1 treatment. In*Trim9* ^+/+^ neurons, bath application of low netrin-1 increased levels of DCC at the plasma membrane surface (Figure 2, p *<* .05), whereas membrane levels of DCC were maintained at baseline at the high concentration.

We previously found that loss of Trim9 was associated with exuberant exocytosis and hyperactivation of DCC (Winkle et al., 2014; Plooster et al., 2017), but we did not examine membrane levels of DCC. *Trim9* ^-/-^ growth cones showed elevated baseline levels of DCC on the membrane compared to *Trim9* ^+/+^ growth cones (Figure 2C, p*<*0.03). DCC was decreased with low netrin-1 treatment compared to sham control in *Trim9* ^-/-^ growth cones (p*<*0.05), whereas surface levels of DCC remained unchanged with high netrin-1treatment (Figure 2C). Thus, TRIM9 deletion leads to the deregulation of DCC levels at the membrane.

Unfortunately, we were unable to identify an antibody that recognizes an extracellular epitope on UNC5C to perform parallel measurements of UNC5C levels at the surface of the neuron. We used an antibody that recognizes the cytoplasmic tail of UNC5C to evaluate total UNC5C levels in the growth cone (Figure 2A). The baseline levels of UNC5C in the growth cone were the same in *Trim9* ^+/+^ and *Trim9* ^-/-^ neurons (Figure 2D). UNC5C levels in the growth cone in *Trim9* ^+/+^ neurons were unaltered by application of a low netrin-1 concentration, but decreased after application of high netrin-1 (Figure 2D, p*<* .05). Levels of UNC5C did not change with the netrin treatment in *Trim9* ^-/-^ growth cones (Figure 2D). Together these data indicate low and high netrin concentrations differentially alter DCC and UNC5C levels in the growth cone, and these responses are TRIM9-dependent.

### TRIM9 regulates membrane levels of DCC and FAK Activity

The published phenotypes in *Trim9* ^-/-^ neurons include exuberant axon branching, exocytosis, and increased growth cone size (Winkle et al., 2014; Menon et al., 2015; Urbina et al., 2018) along with the increased growth cone size and surface levels of DCC reported here (Figure 2). Interestingly these phenocopy *Trim9* ^+/+^ neurons after netrin application. This suggested that Trim9 deletion may induce a state akin to netrin stimulation. Further our previous findings that FAK activation and DCC phosphorylation were aberrantly high in *Trim9* ^-/-^ neurons suggested that “inside-out” activation of DCC occurred in the absence of netrin (Plooster et al., 2017). Neutralizing antibodies that bind the extracellular domain of DCC have been employed to block DCC function (De La Torre et al., 1997; Moore et al., 2008; Uriot et al., 2023). Inspired by these approaches, we used addition of a neutralizing antibody that binds to the extracellular Ig and FN3 domain of DCC to examine if elevated surface levels of DCC on *Trim9* ^-/-^ neurons were associated with increased growth cone area (Figure 3A). Again, we observed that the level of DCC at the cell surface was elevated in *Trim9* ^-/-^ neurons compared to *Trim9* ^+/+^ neurons in the absence of this neutralizing antibody (Figure 3B, p*<*0.0001). Immunostaining with a fluorescent secondary antibody revealed that the neutralizing antibody did bind the neuronal surface (Figure 3A, bottom panel). The DCC neutralizing antibody did not alter growth cone size in *Trim9* ^+/+^ neurons (Figure 3C). However, it did reduce the elevated growth cone area observed in *Trim9* ^-/-^ neurons (Figure 3, p*<*0.0001). These results are consistent with the elevated membrane DCC levels and DCC function in *Trim9* ^-/-^ growth cones causing excessive morphogenesis (Plooster et al., 2017). Several labs have shown that netrin activates the nonreceptor tyrosine kinase FAK, with FAK autophosphorylation (pY397) increasing in the presence of netrin, and subsequently, increased tyrosine phosphorylation of the cytoplasmic tail of DCC Li et al. (2004); Liu et al. (2004); Ren et al. (2004); Plooster et al. (2017). We found that netrin addition also leads to the deubiquitylation of DCC. A non-ubiquitylatable DCC mutant exhibited excessive pY on DCC, suggesting that DCC ubiquitylation maintained DCC in a non-phosphorylated state and, inhibiting aberrant inside-out activation (Plooster et al., 2017). We thus evaluated FAK activity (pY397) upon netrin treatment in *Trim9* ^+/+^ and *Trim9* ^-/-^ cortical neurons in response to attractive and repulsive bath application of murine netrin-1 (Figure 3). In *Trim9* ^+/+^ neurons, FAK activity increased in six of six biological replicates after addition of 600 ng/ml netrin-1, although with the biological variability, this did not reach statistical significant (Figure 2E, p = .15). pY397 did increase significantly after high netrin-1 treatment (p =. 03). Consistent with previous results (Plooster et al., 2017), *Trim9* ^-/-^ cortical neurons had elevated FAK pY397 (p = .008). Further, netrin-1 application did not change pY397 FAK levels in*Trim9* ^-/-^ neurons. These results corroborated our previous findings that netrin-1 increased FAK autophosphorylation, and that *Trim9* ^-/-^ cortical neurons have elevated active FAK signaling, consistent with increased surface levels and inside-out activation of DCC.

**Figure 3:**
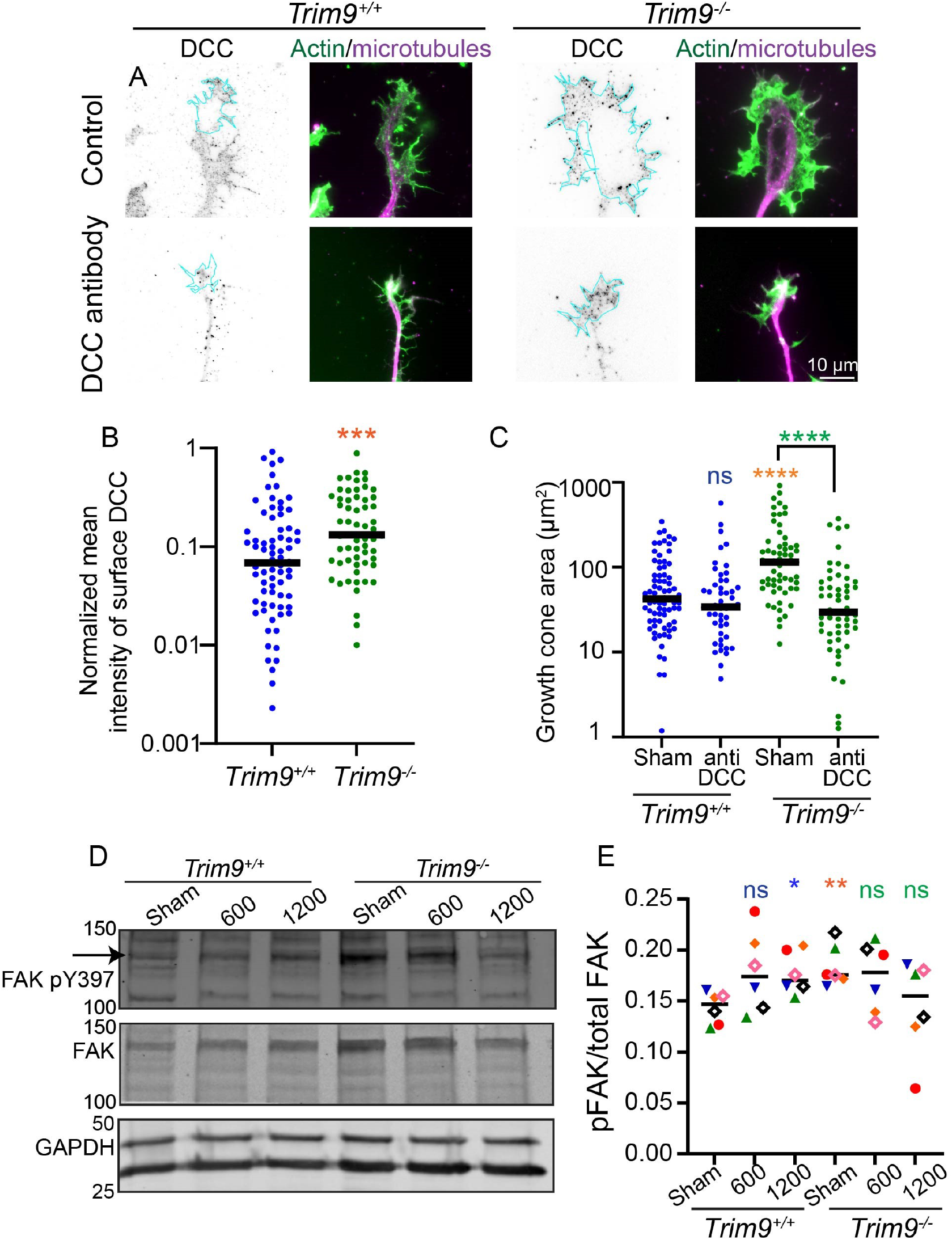
Loss of Trim9 leads to netrin-independent activation of DCC and FAK. **A**. Immunostaining of the extracellular domain of DCC in *Trim9* ^+/+^ and *Trim9* ^-/-^ growth cones with and without addition of neutralizing DCC antibody. In the bottom panel the inverted images show staining with secondary antibody, indicating that neutralizing DCC antibody was bound to the growth cone surface. **B**. Quantification of normalized mean intensity of DCC. **C**. Quantification of growth cone area. More than 50 growth cones in each treatment. Data from three in dependent replicates. The p values were compared by Mann-Whitney test. *** indicates p ¡ 0.001, **** p ¡ 0.0001. **D**. Immunoblot of pY397 (shown with the arrow) and total FAK in *Trim9* ^+/+^ and *Trim9* ^-/-^ neurons, after sham, 600, or 1200 ng/ml netrin-1 application, respectively. **E**. Quantification of pY397FAK/FAK. The p values are from by Mann-Whitney test, using 6 independent biological replicates. * indicates p *<* 0.05, ** p*<*.01. p values from statistical comparisons made between treatment in *Trim9* ^+/+^ samples shown in blue, comparisons between *Trim9* ^+/+^ and and *Trim9* ^-/-^ samples orange, and between *Trim9* ^-/-^ samples are shown in green.

### TRIM9 interaction with DCC and UNC5C

Since Trim9 deletion abrogated repulsive netrin responses, we next asked if TRIM9 colocalized with UNC5C to regulate receptor dynamics and function. Due to issues with specificity of antibodies for endogenous TRIM9 (Winkle et al., 2014; McCormick et al., 2024), and the transient nature of interactions typical of E3 ligases, we used a fluorescently tagged TRIM9 construct that lacks the RING containing ligase domain (TRIM9△RING), known to stabilize protein-protein interactions. This revealed that UNC5C-GFP partially colocalized with TRIM9ΔRING at the tips of dynamic filopodia and near the neuronal surface (Figure 4A, white arrows). To determine if TRIM9 interacted with UNC5C, we used co-immunoprecipitation of epitope-tagged proteins from HEK293 cells, using DCC as a positive control, as its interaction with TRIM9 was reported (Plooster et al., 2017). Consistent with the previous findings, DCC co-immunoprecipitated with TRIM9△RING (Plooster et al, 2017). UNC5C also co-immunoprecipitated with TRIM9△RING, indicating these two proteins may be in a complex. Interestingly, the interaction between TRIM9 and UN5C was seen both in the absence and presence of DCC, indicating that UNC5C interacts with TRIM9 independent of DCC (Figure 4B).

**Figure 4:**
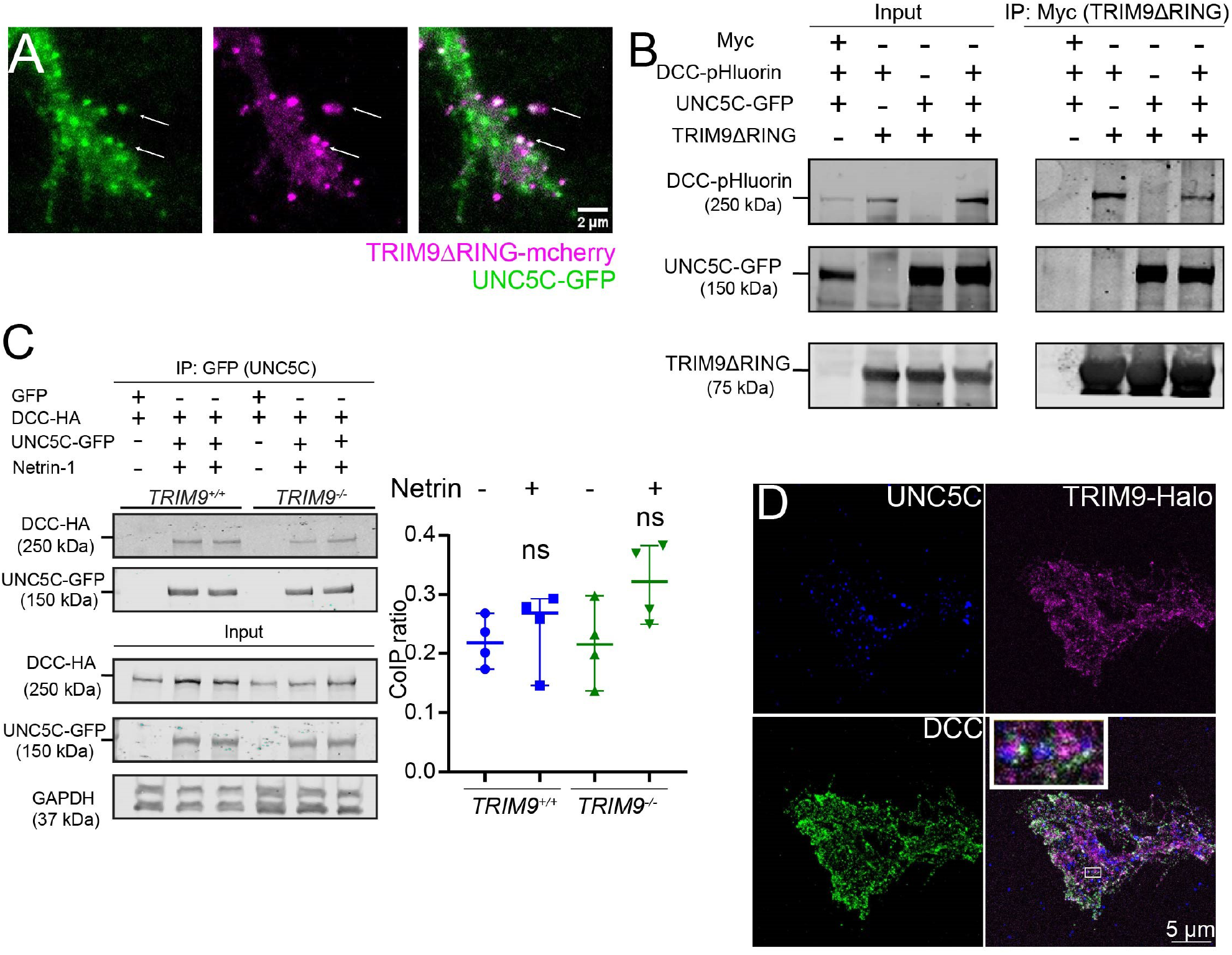
TRIM9 interacts and localizes with DCC and UNC5C. **A**. Representative image of growth cone of *Trim9* ^-/-^ neuron expressing UNC5C-GFP (green) and TRIM9△RING (magenta). Arrows denote colocalization at filopodia tips **B**. Co-IP of DCC-pHluorin and UNC5C-GFP with kennedy in *TRIM9* ^-/-^ HEK293 cells. Representative blot of three replicates. **C**. Co-IP of DCC-HA with UNC5C-GFP in *TRIM9* ^+/+^ and *TRIM9* ^-/-^ HEK293 cells. Data from four independent replicates. Co-IP ratios were compared by unpaired student t test. ns: p *>* 0.05. **D**. Representative example of endogenous DCC, endogenous UNC5C, and TRIM9-Halo colocalization in *Trim9* ^-/-^ growth cone. Inset in merged image shows DCC, UNC5C, and TRIM9 juxtaposition.

These colocalization and coimmunoprecipitation results suggested that DCC, UNC5C, and TRIM9 potentially formed a complex. To determine if UNC5C and DCC interact, which has recently been debated (Keleman and Dickson, 2001; Geisbrecht et al., 2003; Kruger, 2005; Shao et al., 2017), we immunoprecipitated UNC5C-GFP from HEK293 cells. HA-DCC was coimmunoprecipitated with UNC5C, indicating these receptors are in a complex. This interaction was not altered by netrin addition or genetic deletion of TRIM9 (Figure 4C). Thus, biochemical data suggested that DCC and UNC5C were in a complex that is independent of TRIM9 and insensitive to netrin-1 (p *>* 0.05). To evaluate if endogenous DCC and UNC5C receptors colocalized with full length TRIM9, we expressed full length construct of TRIM9 tagged with Halo and used Stimulated Emission Depletion (STED) microscopy on *Trim9* ^-/-^ neurons. This revealed colocalization and juxtaposition of TRIM9-Halo, endogenous DCC, and endogenous UNC5C (Figure 4D).

### Netrin regulates UNC5C dynamics

Netrin-dependent clustering of DCC and UNC5B was reported in HEK293 cells (Gopal et al., 2016, 2017). Our previous work demonstrated netrin-dependent DCC clustering at the plasma membrane in neurons was lost in the absence of Trim9 (Plooster et al., 2017). Whether TRIM9 regulates the surface localization and clustering of UNC5C is not known, as the immunostaining experiments described in Figure 2 only estimated total UNC5C levels. To evaluate UNC5C on the plasma membrane and its regulation by netrin-1 and TRIM9, we generated an UNC5C construct with the pH-sensitive variant of mScarlet (Bindels et al., 2017; Liu et al., 2021) within its extracellular domain (Figure 5A). The fluorescence of pHmScarlet is quenched within acidic vesicles, whereas it fluoresces in the extracellular environment (pH 7.4). Using total internal reflection fluorescence (TIRF) microscopy, we imaged the plasma membrane-localized population of UNC5C-pHmScarlet (Figure 5). We confirmed the membrane localization of UNC5C by altering the pH of vesicles using ammonium chloride, which increased fluorescence (Figure S3A). Intracellular vesicles were observed with increased penetration depth of the evanescent wave, confirming the membrane localization of the UNC5C-pHmScarlet construct in the TIRF illumination mode. We evaluated changes in membrane fluorescence of UNC5C-pHmScarlet upon bath application of 1200 ng/ml netrin-1 (Figure 5A-C). To avoid photobleaching, single images were acquired after 10, 20, 40, and 60 min. Interestingly, fluorescence of surface levels of UNC5C dropped upon netrin-1 application in *Trim9* ^+/+^ neurons. The decrease in was not due to photobleaching, as media sham treated *Trim9* ^+/+^ neurons did not exhibit a drop in fluorescence using the same imaging paradigm (Figure S3B,C). We observed that UNC5C-pHmScarlet fluorescence dropped in a netrin-dependent manner both in *Trim9* ^+/+^ and *Trim9* ^-/-^ cortical neurons (Figure 5A-C). Thus, at these time scales repulsive netrin-1 application reduced surface levels of UNC5C in a TRIM9-independent fashion.

**Figure 5:**
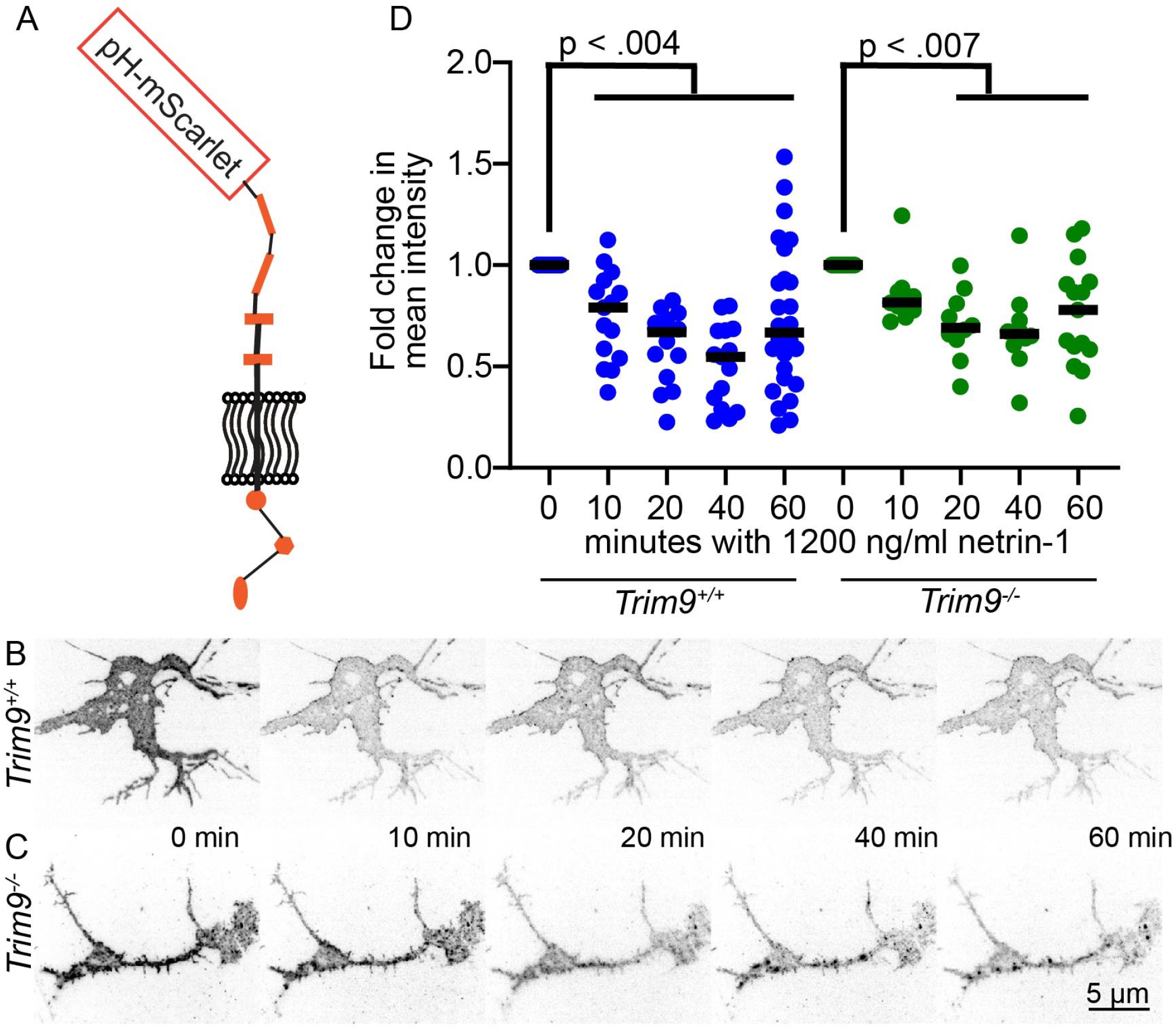
Figure 5. Netrin-1 reduces UNC5C levels on the plasma membrane. **A**. UNC5C construct with pH-mScarlet on the extracellular domain **B-C**. Representative examples of UNC5C-ph-msscarlet expressing *Trim9* ^+/+^ (**B**) *Trim9* ^-/-^ (**C**) cortical neuron. **D**. Fold change in mean intensity after bath application of 1200 ng/ml netrin-1 in *Trim9* ^+/+^ and *Trim9* ^-/-^ cortical neurons. Data from three independent experiments, more than 10 cells per treatment. Means were compared by Kruskal Wallis test with the Dunn’s correction, p values are indicated.

### TRIM9 regulates the mobility of UNC5C in the plasma membrane

Across the surface of the neuron, UNC5C-pHmScarlet fluorescence intensity decreased minutes after bath application of 1200 ng/ml netrin-1 in both *Trim9* ^+/+^ and *Trim9* ^-/-^ neurons ((Figure 5). However, whether TRIM9 or netrin-1 regulated UNC5C mobility at the plasma membrane at faster time scales remained to be seen. To investigate the surface mobility of UNC5C-pHmScarlet, we used fluorescence recovery after photobleaching (FRAP) on the same neuron before and after bath application of 1200 ng/ml netrin-1 ((Figure 6 A,B). The half times (t_2_) of fluorescence recovery, which describe how rapidly bleached molecules leave the bleached region, was measured on same neurons before and after high netrin-1 application. The t_2_ of UNC5C-pHmScarlet in *Trim9* ^+/+^ neurons increased after high netrin-1 application (Figure 6C, p *<* 0.05), indicating netrin-1 slowed mobility. In contrast, there was no change in the t_2_ of UNC5C-pHmScarlet after netrin application in *Trim9* ^+/+^ neurons Figure 6D). However, basally *Trim9* ^-/-^ neurons exhibited a larger t_2_ of UNC5C-pHmScarlet than *Trim9* ^+/+^ neurons (Figure 6E, p *<* 0.05), suggesting that TRIM9 increases UNC5C mobility in the absence of netrin-1. These data indicate netrin-1 and TRIM9 alter UNC5 mobility on the membrane.

**Figure 6:**
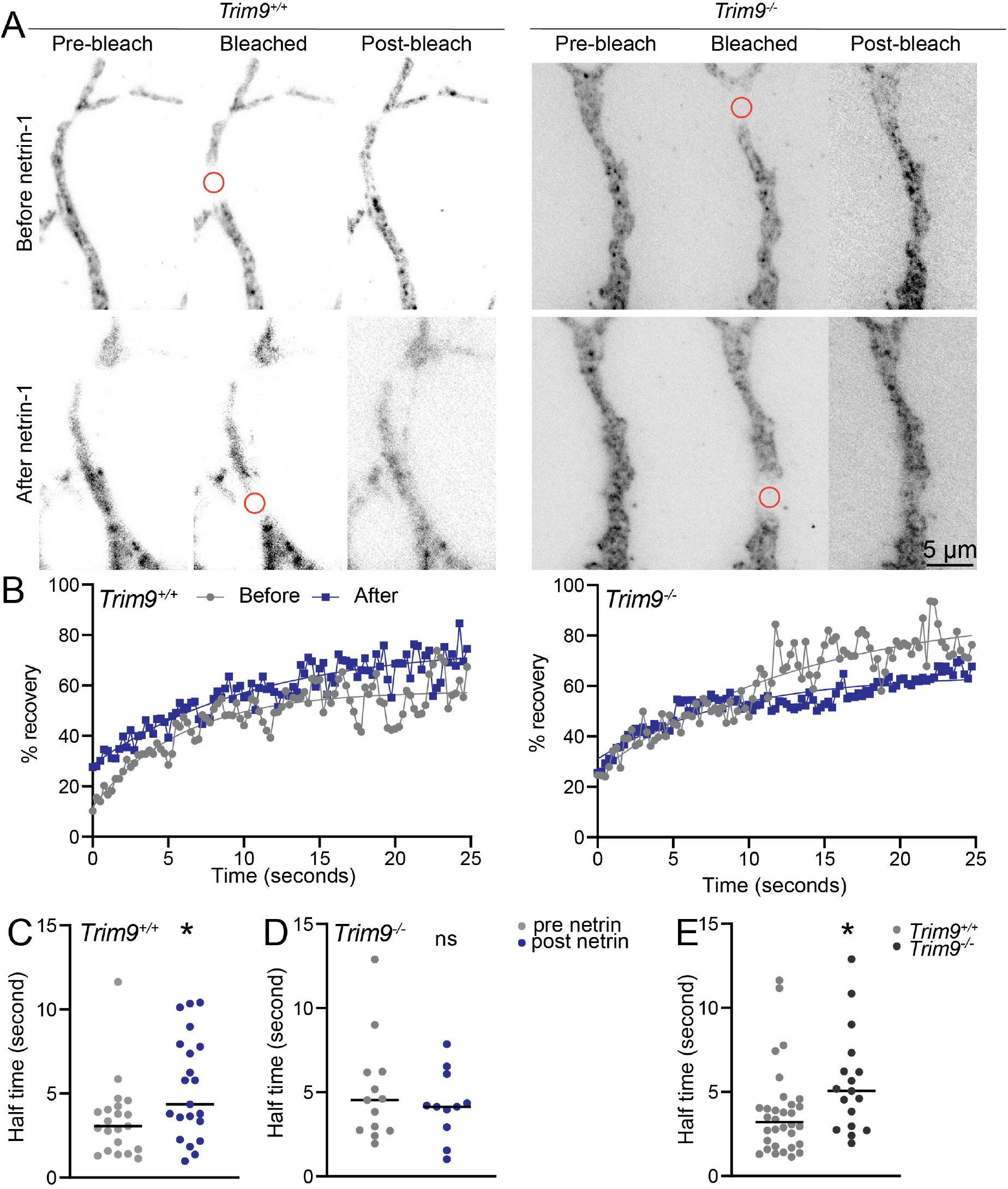
TRIM9 regulates surface mobility of UNC5C. **A**. Representative example of FRAP experiment in *Trim9* ^+/+^ and *Trim9* ^-/-^ cortical neurons, pre and post netrin-1 addition. Red circles denote bleached region of interest. **B**. Representative fluorescence recovery curves fit to single exponential for the examples shown in A. **C-E**. Comparisons of half time of fluorescence recovery. Data from more than three independent replicates. Each category has more than 10 axons. p values are from Wilcoxon test for paired comparisons before and after netrin-1. Half times of recovery in*Trim9* ^+/+^ and *Trim9* ^-/-^ were compared using Mann Whitney test. * indicates p*<* 0.05, ns: p *>* 0.05.

## Discussion

Multiple studies have suggested that netrin-1 can induce both attractive and repulsive responses (Keleman and Dickson, 2001; Fothergill et al., 2014; Shao et al., 2017; Boyer and Gupton, 2018), however, the mechanisms of such responses are not understood. In this study, we identified netrin-1 concentration as a key determinant of neuronal response. The concentration dependent biphasic responses (attractive and repulsive) were observed in response to a gradient of mouse netrin-1, consistent with previously published work using chicken netrin-1 (Menon et al., 2015; Taylor et al., 2015). The conserved guidance responses of mouse and chicken netrin-1 are consistent with the high sequence similarities (88%). Consistent with the biphasic axon turning responses, bath application of low and high concentrations of murine netrin-1 induced attractive and repulsive growth cone responses, respectively. We observed concentration-dependent changes in the levels of netrin receptors DCC and UNC5C: the level of DCC at the plasma membrane increased in response to low netrin, whereas total UNC5C level was reduced in response to high netrin-1. Interestingly, biophysical experiments suggested that DCC has a high affinity for netrin-1, whereas UNC5A, another UNC5 family member that is a netrin receptor, has a lower affinity (Finci et al., 2014). Concentration-dependent changes in netrin responses and receptor levels could be due to distinct affinities of the receptors with netrin-1, which needs to be tested further.

Distinct presentations of netrin-1 provided valuable insights into netrin-dependent responses. For example, we observed repulsive like phenotypes to high netrin-1 concentrations presented in a bath and as a gradient, although these behaviors were different. When presented as an acute bath application of high netrin-1, multiple growth cones underwent collapse or retraction. However, when presented in the form of the gradient, repulsive turning was observed. How netrin-1 is presented in vivo is still debated, but our results suggest growth cone dynamics are influenced by the presentation of netrin-1, which may be a deciding factor in eliciting specific responses.

### TRIM9 is a novel regulator of netrin-1 biphasic responses

Using multiple netrin-1 presentations, we showed that Trim9 deletion impairs both attractive and repulsive netrin-1 responses. Further at baseline, the phenotypes of *Trim9* ^-/-^ neurons often phenocopied netrin addition. For example, there was a baseline increase in growth cone area, FAK activation, levels of DCC on the plasma membrane, and slowing of UNC5C mobility at the plasma membrane in*Trim9* ^-/-^ neurons that was similar to phenotypes observed in Trim9+/+ neurons following application of netrin-1. Netrin-dependent membrane recruitment and clustering of DCC has been shown previously (Gopal et al., 2016, 2017; Plooster et al., 2017). Using a neutralizing antibody that binds to an extracellular epitope in the netrin-binding region of DCC reduced growth cone area of *Trim9* ^-/-^ growth cones. Together, along with our previous study showing TRIM9-dependent nondegradative ubiquitination of DCC (Plooster et al., 2017), these data suggest that TRIM9 acts as a negative cytoplasmic regulator of DCC “inside-out” activation in the absence of netrin-1, and thus is necessary for netrin-mediated activation.

### Regulation of FAK activity in netrin-1 responses

In an attractive netrin-1 response, activation of FAK and subsequent phosphorylation of the cytoplasmic tail of DCC stimulate downstream signaling and morphogenesis (Li et al., 2004; Ren et al., 2004; Moore et al., 2012; Plooster et al., 2017; Mahadik and Lundquist, 2023). In mouse cortical neurons and Xenopus retinal ganglion cells, autophosphorylation of FAK in response to netrin-1 application recruits Src kinase and multiple cytoskeletal regulators, including the Arp2/3 complex and consequentially, actin polymerization takes place (Li et al., 2004; Liu et al., 2004; Meriane et al., 2004). Multiple cell biological studies have focused on these signaling pathways and morphological responses in response to presumably attractive concentrations of netrin-1 (Table S1). We also observed elevated FAK activity (pY397) upon bath application of high concentrations of netrin-1 in *Trim9* ^+/+^ neurons, suggesting a role of FAK signaling in repulsive responses. However, how attractive versus repulsive responses are distinguished downstream of FAK autophosphorylation remains a conundrum. Repulsive cues like semaphorin A also increase pY397 levels in mouse cortico-hippocampal neurons, whereas phosphorylation of other tyrosine residues on FAK decreases, suggesting that pY397 magnitude and/or downstream FAK phos-phorylation status could provide an additional layer of the regulation in guidance responses (**?**Navarro and Rico, 2014).

We observed elevated basal FAK activity (pY397) in *Trim9* ^-/-^ neurons, which remained unchanged with netrin-1 application, highlighting regulation of FAK activity involves TRIM9. Based on super-resolution imaging, TRIM9 colocalizes with endogenous DCC and UNC5C, highlighting its proximity for negatively regulating FAK activity and inside-out signaling at receptor clusters. Particularly, in the absence of netrin-1, negative regulation of FAK activity is critical to maintain the cell shape (Plooster et al., 2017) and prime the neuron to respond to netrin-1. Upon stimulation, negative regulation of FAK by TRIM9 must be relieved to facilitate FAK activation and downstream signaling at the receptor clusters.

### Functions of TRIM9 in netrin-dependent repulsion

One interesting in vivo phenotype associated with murine Trim9 deletion is increased cortical axon projection to the internal capsule (Menon et al., 2015). Interestingly, Unc5c mutants also exhibit this guidance defect, in which netrin-sensitive axonal tracts are misrouted (Srivatsa et al., 2014). Here we find that TRIM9 regulates the mobility of UNC5C within the plasma membrane. There are multiple potential mechanisms by which TRIM9 may regulate UNC5C mobility, including by regulating membrane composition, the cortical actin cytoskeleton, or interaction with other receptors. Previous work found that cholesterol-enriched lipid rafts are critical for the localization of UNC5C receptors and netrin-dependent repulsion (Hernaiz-Llorens et al., 2021). TRIM9 regulates SNARE complex formation and membrane expansion (Winkle et al., 2014); this may impact plasma membrane composition and thus, UNC5C mobility. Our recent interactome study (Menon et al., 2021) highlights multiple actin binding proteins as TRIM9 interactors including Ena/VASP proteins and Myo16 (Menon et al., 2015) (Menon et al., 2015). These interactions poise TRIM9 to regulate cortical actin and hence mobility of UNC5C and other receptors. If so, the slower recovery after photobleaching of UNC5C-pHmScarlet in *Trim9* ^-/-^ neurons may be due to an altered cortical cytoskeletal network. Alternatively, TRIM9 may alter UNC5C interactions with other transmembrane proteins. Surprisingly we did not observe netrin or TRIM9-sensitive interactions between UNC5C and DCC, even though we have previously seen TRIM9 is critical for netrin-dependent clustering of DCC (Plooster et al., 2017). Whether TRIM9 alters UNC5C interaction with other receptors like DSCAM is not known.

### Netrin dependent mechanisms of UNC5C regulation on the membrane

We observed that repulsive concentrations of netrin-1 reduced the amount of UNC5C on the neuronal surface in a TRIM9-independent fashion. Although the mechanisms by which UNC5C removal occurs is not known, there are several possibilities. Growth cone collapse requires removal of plasma membrane (Fournier et al., 2000; Tojima et al., 2010). Endocytosis of membrane material and UNC5C could accomplish both membrane reduction and UNC5C removal. Previously, PKC-dependent phos-phorylation of UNC5A was suggested to induce endocytosis of UNC5A in mouse granule neurons (Bartoe et al., 2006). Whether the decrease in UNC5C membrane fluorescence reported here is due to PKC-dependent endocytosis is not known. Alternatively, other extracellular mechanisms like protease-dependent receptor shedding (Gatto et al., 2014; Van Erp et al., 2015) may remove UNC5C in response to repulsive netrin-1. Interestingly, DCC is a substrate for protease-mediated ectodomain shedding. Inhibition of proteolytic activity caused an increase in DCC protein levels (Galko and Tessier-Lavigne, 2000). Whether UNC5C undergoes receptor shedding and how that influences repulsion is not known. Interestingly, UNC5C also interacts with the exocytic t-SNARE protein syntaxin-1. Syntaxin-1 was shown to be involved in netrin dependent repulsion via a role in micropinocytosis (Martínez-Mármol et al., 2023). Syntaxin-1 and UNC5C receptors were shown to form membrane nanoclusters (Bademosi et al., 2016; Hernaiz-Llorens et al., 2021). These structures could potentially initiate membrane internalization during repulsion. These or other mechanisms or combination of them (shedding, endocytosis and micropinocytosis) are likely to UNC5 regulate receptor levels and membrane recycling during repulsion.

## Supporting information

Video3: Bath application of netrin-1 induces growth cone retraction. An example of growing Trim9+/+ axon that undergoes retraction upon bath applicati

Video 4: Example of a paused growth cone transitioning to ruffling. An example of a paused Trim9+/+ axon that undergoes dynamic ruffling upon bath app

Video1: Attractive axon turning in a low netrin-1 gradient. An example of a Trim9+/+ axon in the low end of the netrin-1 gradient turning up toward th

Video2: Repulsive axon turning in a high netrin-1 gradient. An example of Trim9+/+ axons in the high end of the netrin-1 gradient turning down away fr

## Acknowledgements

We thank Shalini Menon and Nicholas Boyer for performing preliminary axon turning and growth cone experiments using chicken netrin-1. We thank Caroline Monkiewicz, Shalini Menon, and Joy Furigay for performing initial TRIM9:UNC5C co-immunopreciptiation experiments. Together these experiments motivated this study. We thank Samantha Butler for excellent conversation and generous sharing of murine netrin-1 plasmids. We thank Elliot Evans and Aneri Shah for their contributions to the project, David Adalsteinsson for the generous help with the ImageTank software. We acknowledge the Hooker Imaging Core (UNC) for facilitating STED imaging and Wendy Salmon for the technical assistance. We thank Janee Cadlet and Natalia Riddick for mouse colony management assistance. This work was supported by NIH R35GM135160 (S.L.G), R21NS132364 (S.L.G), and NIH S10OD030300 (S.L.G.).

---

**Figure S1.**
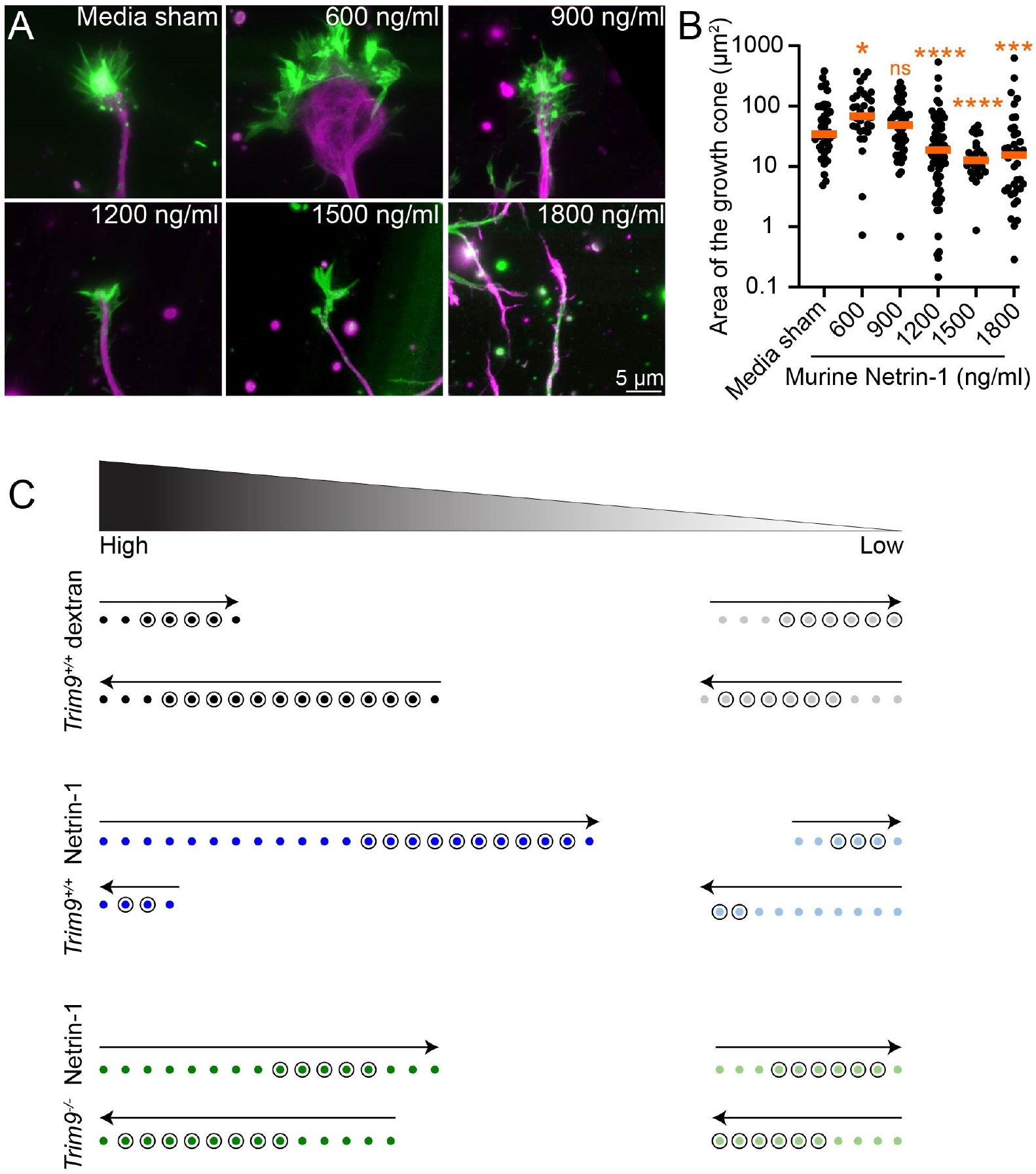
Biphasic growth cone responses to murine netrin-1 treatment. **A**. Immunostaining of *Trim9*^*+/+*^ growth cones treated with indicated concentrations of murine netrin-1. F-actin is stained with phalloidin (green) and microtubules (magenta) with *β*-III tubulin antibody. **B** Quantification of growth cone area. Data from two experiments, more than 25 growth cones in each treatment. The *p* (*<*0.0001) value was compared by Kruskal-Wallis test (multiple comparisons were corrected by Dunn’s test). **C**. Summary of final growth direction of axons (arrows) in microfluidic devices of indicated genotypes within high and low regions of a dextran or netrin-1 gradient. Circled dots indicate axons that turned *<*15° during the analysis time, and thus showed low turning angles in Figure 1C.

**Figure S2.**
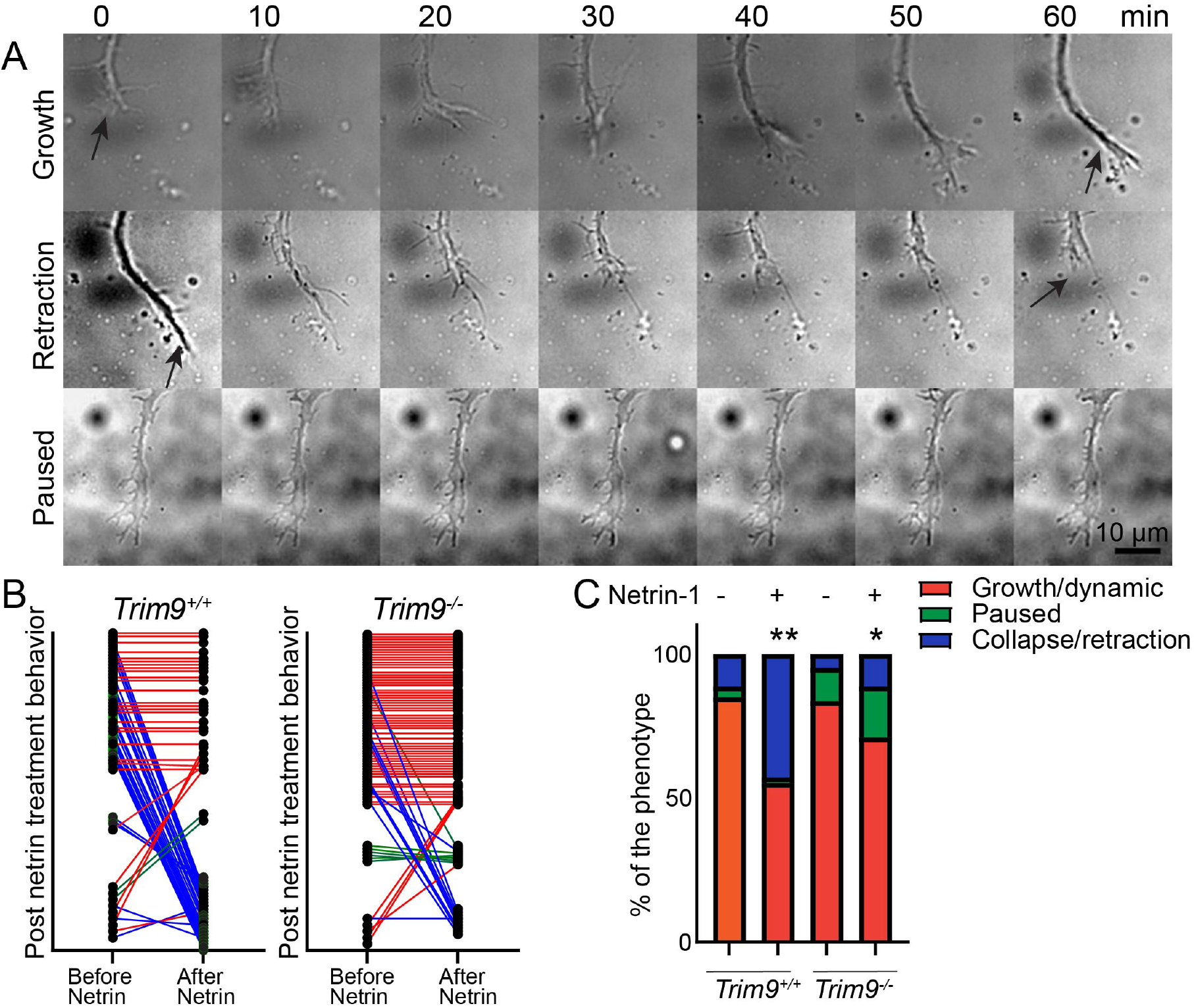
Netrin-dependent growth cone dynamics are altered in *Trim9*^*-/-*^ neurons. **A**. Examples of growing (+) (before netrin), retracting axon (-) (after netrin) and paused axon (before netrin) (=) of *Trim9*^*+/+*^ cortical neurons. **B**. Growth cone response of individual growth cones are shown. Orange, blue, and green line indicate growing, retracting and paused axons post netrin treatment. **C**. Percentage of growth cones in each behavior before and after netrin addition. Data from three replicates (n=62, n=86 for *Trim9*^*+/+*^ and *Trim9*^*-/-*^, respectively). Statistical comparision using *χ*^2^ test with Bonferroni correction. * p *<* .05, ** p*<* .01. Please refer to Video 3 and 4.

**Figure S3.**
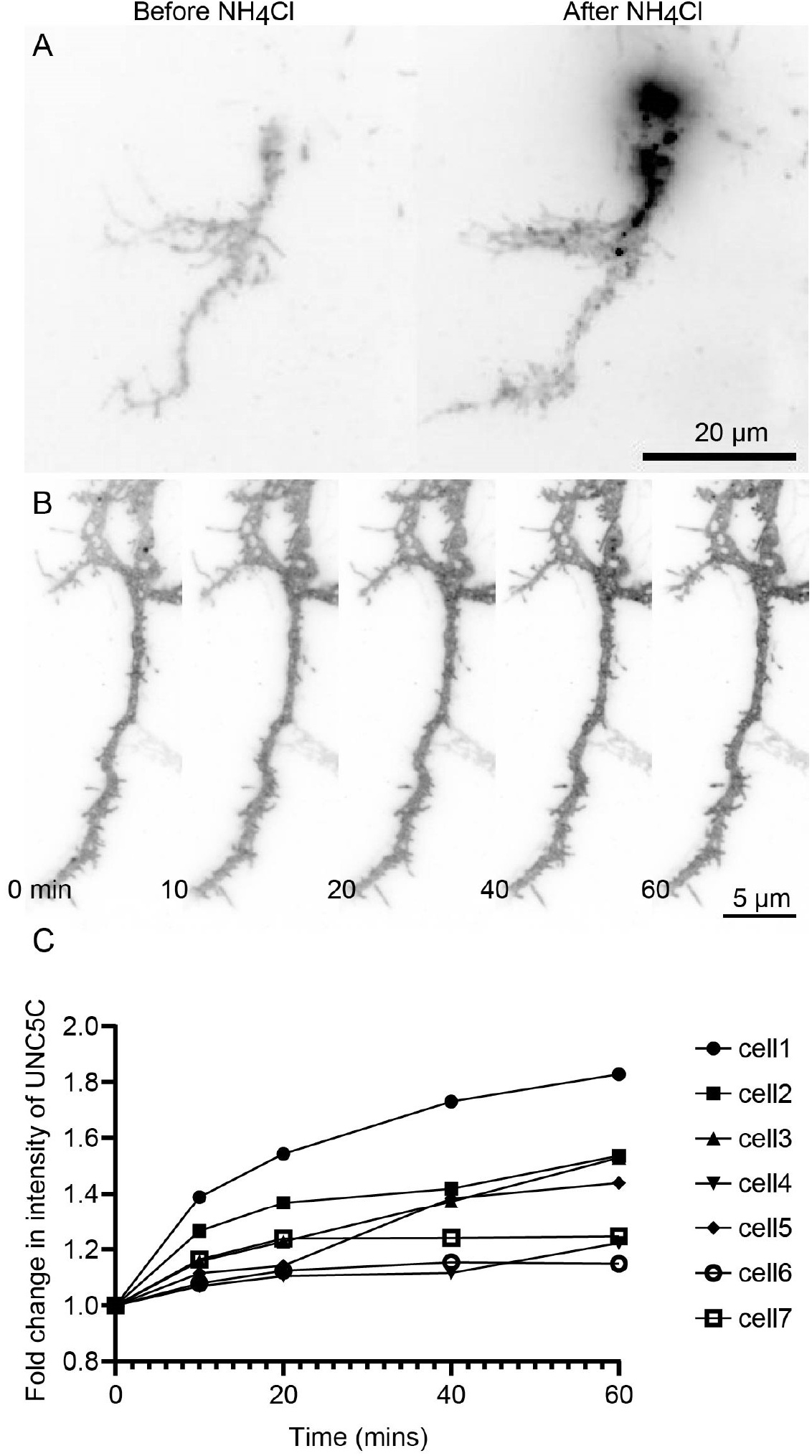
Validation of UNC5C-pHmScarlet membrane localization. **A**. An example of a neuron expressing UNC5C-pHmScarlet before and after addition of ammonium chloride (100 μM) imaged in TIRF mode. **B**. An example of UNC5C-pHmScarlet expressing neuron. **C**. Fold change in mean intensity with the addition media is shown. Media was added at 10 min.

## Supplemental Video Legends

Video1: **Attractive axon turning in a low netrin-1 gradient**. An example of a *Trim9*^*+/+*^, axon in the low end of the netrin-1 gradient turning up toward the netrin source (attractive response). Direction of gradient indicated by triangle. Elapsed time in minutes.

**Video2: Repulsive axon turning in a high netrin-1 gradient**. An example of *Trim9*^*+/+*^, axons in the high end of the netrin-1 gradient turning down away from the netrin source (repulsive response). Direction of gradient indicated by triangle. Elapsed time in minutes.

**Video3: Bath application of netrin-1 induces growth cone retraction**. An example of growing *Trim9*^*+/+*^,axon that undergoes retraction upon bath application of 1200 ng/ml netrin-1 addition. Netrin-1 was added at 1 hr. Elapsed time in hr:mins.

**Video 4:** Example of a paused growth cone transitioning to ruffling. An example of a paused *Trim9*^*+/+*^, axon that undergoes dynamic ruffling upon bath application of 1200 ng/ml netrin-1 addition. Netrin-1 was added at 1 hr. Elapsed time in hr:mins.

## References

Ackerman, S. L., Kozak, L. P., Przyborski, S. A., Rund, L. A., Boyer, B. B., and Knowles, B. B. (1997). The mouse rostral cerebellar malformation gene encodes an unc-5-like protein. Nature, 386(6627):838–842.

Bademosi, A. T., Lauwers, E., Padmanabhan, P., Odierna, L., Chai, Y. J., Papadopulos, A., Goodhill, G. J., Verstreken, P., Van Swinderen, B., and Meunier, F. A. (2016). In vivo single-molecule imaging of syntaxin1a reveals polyphosphoinositide-and activity-dependent trapping in presynaptic nanoclusters. Nature communications, 7(1):13660.

Bartoe, J. L., McKenna, W. L., Quan, T. K., Stafford, B. K., Moore, J. A., Xia, J., Takamiya, K., Huganir, R. L., and Hinck, L. (2006). Protein interacting with c-kinase 1/protein kinase cα-mediated endocytosis converts netrin-1-mediated repulsion to attraction. Journal of Neuroscience, 26(12):3192–3205.

Bin, J. M., Han, D., Sun, K. L. W., Croteau, L.-P., Dumontier, E., Cloutier, J.-F., Kania, A., and Kennedy, T. E. (2015). Complete loss of netrin-1 results in embryonic lethality and severe axon guidance defects without increased neural cell death. Cell reports, 12(7):1099–1106.

Bindels, D. S., Haarbosch, L., Van Weeren, L., Postma, M., Wiese, K. E., Mastop, M., Aumonier, S., Gotthard, G., Royant, A., Hink, M. A., et al. (2017). mscarlet: a bright monomeric red fluorescent protein for cellular imaging. Nature methods, 14(1):53–56.

Boyer, N. P. and Gupton, S. L. (2018). Revisiting netrin-1: one who guides (axons). Frontiers in cellular neuroscience, 12:221.

Boyer, N. P., McCormick, L. E., Menon, S., Urbina, F. L., and Gupton, S. L. (2019). A pair of e3 ubiquitin ligases compete to regulate filopodial dynamics and axon guidance. Journal of Cell Biology, 219(1):e201902088.

Bradford, D., Cole, S. J., and Cooper, H. M. (2009). Netrin-1: diversity in development. The international journal of biochemistry & cell biology, 41(3):487–493.

Braisted, J. E., Catalano, S. M., Stimac, R., Kennedy, T. E., Tessier-Lavigne, M., Shatz, C. J., and O’Leary, D. D. (2000). Netrin-1 promotes thalamic axon growth and is required for proper development of the thalamocortical projection. Journal of Neuroscience, 20(15):5792–5801.

Dailey-Krempel, B., Martin, A. L., Jo, H.-N., Junge, H. J., and Chen, Z. (2023). A tug of war between dcc and robo1 signaling during commissural axon guidance. Cell reports, 42(5).

Day, C. A., Kraft, L. J., Kang, M., and Kenworthy, A. K. (2012). Analysis of protein and lipid dynamics using confocal fluorescence recovery after photobleaching (frap). Current protocols in cytometry, 62(1):2–19.

De La Torre, J. R., Höpker, V. H., Ming, G.-l., Poo, M.-m., Tessier-Lavigne, M., Hemmati-Brivanlou, A., and Holt, C. E. (1997). Turning of retinal growth cones in a netrin-1 gradient mediated by the netrin receptor dcc. Neuron, 19(6):1211–1224.

Degasperi, A., Birtwistle, M. R., Volinsky, N., Rauch, J., Kolch, W., and Kholodenko, B. N. (2014). Evaluating strategies to normalise biological replicates of western blot data. PloS one, 9(1):e87293.

Finci, L. I., Krüger, N., Sun, X., Zhang, J., Chegkazi, M., Wu, Y., Schenk, G., Mertens, H. D., Svergun, D. I., Zhang, Y., et al. (2014). The crystal structure of netrin-1 in complex with dcc reveals the bifunctionality of netrin-1 as a guidance cue. Neuron, 83(4):839–849.

Fothergill, T., Donahoo, A.-L. S., Douglass, A., Zalucki, O., Yuan, J., Shu, T., Goodhill, G. J., and Richards, L. J. (2014). Netrin-dcc signaling regulates corpus callosum formation through attraction of pioneering axons and by modulating slit2-mediated repulsion. Cerebral cortex, 24(5):1138–1151.

Fournier, A. E., Nakamura, F., Kawamoto, S., Goshima, Y., Kalb, R. G., and Strittmatter, S. M. (2000). Semaphorin3a enhances endocytosis at sites of receptor–f-actin colocalization during growth cone collapse. The Journal of cell biology, 149(2):411–422.

Galko, M. J. and Tessier-Lavigne, M. (2000). Function of an axonal chemoattractant modulated by metalloprotease activity. Science, 289(5483):1365–1367.

Gatto, G., Morales, D., Kania, A., and Klein, R. (2014). Epha4 receptor shedding regulates spinal motor axon guidance. Current Biology, 24(20):2355–2365.

Geisbrecht, B. V., Dowd, K. A., Barfield, R. W., Longo, P. A., and Leahy, D. J. (2003). Netrin binds discrete subdomains of dcc and unc5 and mediates interactions between dcc and heparin. Journal of Biological Chemistry, 278(35):32561–32568.

Goldowitz, D., Hamre, K. M., Przyborski, S. A., and Ackerman, S. L. (2000). Granule cells and cerebellar boundaries: Analysis ofunc5h3 mutant chimeras. Journal of Neuroscience, 20(11):4129–4137.

Gopal, A. A., Rappaz, B., Rouger, V., Martyn, I. B., Dahlberg, P. D., Meland, R. J., Beamish, I. V., Kennedy, T. E., and Wiseman, P. W. (2016). Netrin-1-regulated distribution of unc5b and dcc in live cells revealed by ticcs. Biophysical Journal, 110(3):623–634.

Gopal, A. A., Ricoult, S. G., Harris, S. N., Juncker, D., Kennedy, T. E., and Wiseman, P. W. (2017). Spatially selective dissection of signal transduction in neurons grown on netrin-1 printed nanoarrays via segmented fluorescence fluctuation analysis. ACS nano, 11(8):8131–8143.

Hernaiz-Llorens, M., Roselló-Busquets, C., Durisic, N., Filip, A., Ulloa, F., Martínez-Mármol, R., and Soriano, E. (2021). Growth cone repulsion to netrin-1 depends on lipid raft microdomains enriched in unc5 receptors. Cellular and Molecular Life Sciences, 78:2797–2820.

Keino-Masu, K., Masu, M., Hinck, L., Leonardo, E. D., Chan, S. S.-Y., Culotti, J. G., and Tessier-Lavigne, M. (1996). Deleted in colorectal cancer (dcc) encodes a netrin receptor. Cell, 87(2):175–185.

Keleman, K. and Dickson, B. J. (2001). Short-and long-range repulsion by the drosophila unc5 netrin receptor. Neuron, 32(4):605–617.

Kennedy, T. E., Serafini, T., De La Torre, J., and Tessier-Lavigne, M. (1994). Netrins are diffusible chemotropic factors for commissural axons in the embryonic spinal cord. Cell, 78(3):425–435.

Kim, D. and Ackerman, S. L. (2011). The unc5c netrin receptor regulates dorsal guidance of mouse hindbrain axons. Journal of Neuroscience, 31(6):2167–2179.

Kruger, R. P. (2005). Mechanisms of signaling for netrin and its receptors DCC and Unc5. University of Michigan.

Leonardo, E. D., Hinck, L., Masu, M., Keino-Masu, K., Ackerman, S. L., and Tessier-Lavigne, M. (1997). Vertebrate homologues of c. elegans unc-5 are candidate netrin receptors. Nature, 386(6627):833–838.

Li, W., Lee, J., Vikis, H. G., Lee, S.-H., Liu, G., Aurandt, J., Shen, T.-L., Fearon, E. R., Guan, J.-L., Han, M., et al. (2004). Activation of fak and src are receptor-proximal events required for netrin signaling. Nature neuroscience, 7(11):1213–1221.

Liu, A., Huang, X., He, W., Xue, F., Yang, Y., Liu, J., Chen, L., Yuan, L., and Xu, P. (2021). phmscarlet is a ph-sensitive red fluorescent protein to monitor exocytosis docking and fusion steps. Nature Communications, 12(1):1413.

Liu, G., Beggs, H., Jürgensen, C., Park, H.-T., Tang, H., Gorski, J., Jones, K. R., Reichardt, L. F., Wu, J., and Rao, Y. (2004). Netrin requires focal adhesion kinase and src family kinases for axon outgrowth and attraction. Nature neuroscience, 7(11):1222–1232.

Liu, G., Li, W., Wang, L., Kar, A., Guan, K.-L., Rao, Y., and Wu, J. Y. (2009). Dscam functions as a netrin receptor in commissural axon pathfinding. Proceedings of the National Academy of Sciences, 106(8):2951– 2956.

Mahadik, S. S. and Lundquist, E. A. (2023). A short isoform of the unc-6/netrin receptor unc-5 is required for growth cone polarity and robust growth cone protrusion in caenorhabditis elegans. Frontiers in Cell and Developmental Biology, 11.

Martínez-Mármol, R., Muhaisen, A., Cotrufo, T., Roselló-Busquets, C., Ros, O., Hernaiz-Llorens, M., Pérez-Branguli, F., Andrés, R. M., Parcerisas, A., Pascual, M., et al. (2023). Syntaxin-1 is necessary for unc5a-c/netrin-1-dependent macropinocytosis and chemorepulsion. Frontiers in Molecular Neuroscience, 16.

McCormick, L. E., Evans, E. B., Barker, N. K., Herring, L. E., Diering, G. H., and Gupton, S. L. (2024). The e3 ubiquitin ligase trim9 regulates synaptic function and actin dynamics in response to netrin-1. Molecular Biology of the Cell, 35(5):ar67.

Menon, S., Boyer, N. P., Winkle, C. C., McClain, L. M., Hanlin, C. C., Pandey, D., Rothenfußer, S., Taylor, A. M., and Gupton, S. L. (2015). The e3 ubiquitin ligase trim9 is a filopodia off switch required for netrin-dependent axon guidance. Developmental cell, 35(6):698–712.

Menon, S., Goldfarb, D., Ho, C. T., Cloer, E. W., Boyer, N. P., Hardie, C., Bock, A. J., Johnson, E. C., Anil, J., Major, M. B., et al. (2021). The trim9/trim67 neuronal interactome reveals novel activators of morphogenesis. Molecular biology of the cell, 32(4):314–330.

Meriane, M., Tcherkezian, J., Webber, C. A., Danek, E. I., Triki, I., McFarlane, S., Bloch-Gallego, E., and Lamarche-Vane, N. (2004). Phosphorylation of dcc by fyn mediates netrin-1 signaling in growth cone guidance. The Journal of cell biology, 167(4):687–698.

Mitchell, K. J., Doyle, J. L., Serafini, T., Kennedy, T. E., Tessier-Lavigne, M., Goodman, C. S., and Dickson, B. J. (1996). Genetic analysis of netrin genes in drosophila: Netrins guide cns commissural axons and peripheral motor axons. Neuron, 17(2):203–215.

Moore, S. W., Correia, J. P., Lai Wing Sun, K., Pool, M., Fournier, A. E., and Kennedy, T. E. (2008). Rho inhibition recruits dcc to the neuronal plasma membrane and enhances axon chemoattraction to netrin 1.

Moore, S. W., Tessier-Lavigne, M., and Kennedy, T. E. (2007). Netrins and their receptors. Axon Growth and Guidance, pages 17–31.

Moore, S. W., Zhang, X., Lynch, C. D., and Sheetz, M. P. (2012). Netrin-1 attracts axons through fak-dependent mechanotransduction. Journal of Neuroscience, 32(34):11574–11585.

Navarro, A. I. and Rico, B. (2014). Focal adhesion kinase function in neuronal development. Current opinion in neurobiology, 27:89–95.

O’Shaughnessy, E. C., Lam, M., Ryken, S. E., Wiesner, T., Lukasik, K., Zuchero, J. B., Leterrier, C., Adalsteinsson, D., and Gupton, S. L. (2024). phusion: A robust and versatile toolset for automated detection and analysis of exocytosis. Journal of Cell Science, pages jcs–261828.

O’Shaughnessy, E. C., Stone, O. J., LaFosse, P. K., Azoitei, M. L., Tsygankov, D., Heddleston, J. M., Legant, W. R., Wittchen, E. S., Burridge, K., Elston, T. C., et al. (2019). Software for lattice light-sheet imaging of fret biosensors, illustrated with a new rap1 biosensor. Journal of Cell Biology, 218(9):3153–3160.

Phan, K. D., Croteau, L.-P., Kam, J. W. K., Kania, A., Cloutier, J.-F., and Butler, S. J. (2011). Neogenin may functionally substitute for dcc in chicken. PloS one, 6(7):e22072.

Plooster, M., Menon, S., Winkle, C. C., Urbina, F. L., Monkiewicz, C., Phend, K. D., Weinberg, R. J., and Gupton, S. L. (2017). Trim9-dependent ubiquitination of dcc constrains kinase signaling, exocytosis, and axon branching. Molecular biology of the cell, 28(18):2374– 2385.

Powell, A. W., Sassa, T., Wu, Y., Tessier-Lavigne, M., and Polleux, F. (2008). Topography of thalamic projections requires attractive and repulsive functions of netrin-1 in the ventral telencephalon. PLoS biology, 6(5):e116.

Purohit, A. A., Li, W., Qu, C., Dwyer, T., Shao, Q., Guan, K.-L., and Liu, G. (2012). Down syndrome cell adhesion molecule (dscam) associates with uncoordinated-5c (unc5c) in netrin-1-mediated growth cone collapse. Journal of Biological Chemistry, 287(32):27126–27138.

Rajasekharan, S. and Kennedy, T. E. (2009). The netrin protein family. Genome biology, 10:1–8.

Ren, X.-r., Ming, G.-l., Xie, Y., Hong, Y., Sun, D.-m., Zhao, Z.-q., Feng, Z., Wang, Q., Shim, S., Chen, Z.-f., et al. (2004). Focal adhesion kinase in netrin-1 signaling. Nature neuroscience, 7(11):1204–1212.

Serafini, T., Colamarino, S. A., Leonardo, E. D., Wang, H., Beddington, R., Skarnes, W. C., and Tessier-Lavigne, M. (1996). Netrin-1 is required for commissural axon guidance in the developing vertebrate nervous system. Cell, 87(6):1001–1014.

Shao, Q., Yang, T., Huang, H., Alarmanazi, F., and Liu, G. (2017). Uncoupling of unc5c with polymerized tubb3 in microtubules mediates netrin-1 repulsion. Journal of Neuroscience, 37(23):5620–5633.

Srivatsa, S., Parthasarathy, S., Britanova, O., Bormuth, I., Donahoo, A.-L., Ackerman, S. L., Richards, L. J., and Tarabykin, V. (2014). Unc5c and dcc act down-stream of ctip2 and satb2 and contribute to corpus callosum formation. Nature communications, 5(1):3708.

Taylor, A., Menon, S., and Gupton, S. (2015). Passive microfluidic chamber for long-term imaging of axon guidance in response to soluble gradients. Lab on a Chip, 15(13):2781–2789.

Tojima, T., Itofusa, R., and Kamiguchi, H. (2010). Asymmetric clathrin-mediated endocytosis drives repulsive growth cone guidance. Neuron, 66(3):370–377.

Urbina, F. L., Gomez, S. M., and Gupton, S. L. (2018). Spatiotemporal organization of exocytosis emerges during neuronal shape change. Journal of Cell Biology, 217(3):1113–1128.

Uriot, K., Blanc, O., Audugé, N., Faklaris, O., Chaverot, N., Bloch-Gallego, E., Borghi, N., Coppey-Moisan, M., and Girard, P. P. (2023). Interactions of netrin-1 through its glycosylation sites immobilize deleted in colorectal cancer (dcc) by favoring its constitutive clustering. bioRxiv, pages 2023–10.

Van Erp, S., van den Heuvel, D. M., Fujita, Y., Robinson, R. A., Hellemons, A. J., Adolfs, Y., Van Battum, E. Y., Blokhuis, A. M., Kuijpers, M., Demmers, J. A., et al. (2015). Lrig2 negatively regulates ectodomain shedding of axon guidance receptors by adam proteases. Developmental cell, 35(5):537–552.

Winkle, C. C., McClain, L. M., Valtschanoff, J. G., Park, C. S., Maglione, C., and Gupton, S. L. (2014). A novel netrin-1–sensitive mechanism promotes local snare-mediated exocytosis during axon branching. Journal of Cell Biology, 205(2):217–232.

