## Supplementary material for "TRIM9 controls growth cone responses to netrin through DCC and UNC5C": Table S1

| References | Netrin source | Concentration | Incubation | Neuronal cell type | Results |
|---|---|---|---|---|---|
| (Li *et al.*, 2004) | Chicken-myc netrin from HEK293 stable cell lines | 2.5 ug of netrin gradient | one hour exposure of a gradient using micropipette | *X. laevis* spinal neurons | Netrin dependent attractive response was shown using explant culture and micropipette assay. Netrin addition activated FAK and Src and further stimulated tyrosine phosphorylation of DCC in an attractive response. |
| (Liu *et al.*, 2004) | Chicken netrin/ netrin conditioned media | 100-500 ng/ml of purified netrin | 16-18 hour for explant culture, 5-60 mins for FAK phosphorylation | spinal and cortical neurons rat and mice | Netrin induced attraction required FAK and SRC kinase activity. |
| (Ren *et al.*, 2004) | Human netrin-1 | 200 ng/ml netrin | 5-60 mins for FAK phosphorylation assay | Rat cortical neurons | Netrin-1 induced FAK and DCC tyrosin phosphorylation |
| (Moore *et al.*, 2012) | Chicken netrin | 200 ng/ml netrin with 2 ug/ml heparin coated on poly-L-lysine plates and in collagen gels. | 16 hours in collagen gels | Mouse cortical neurons | Netrin-1 can regulate FAK dependent mechanotransduction. |
| (Plooster *et al.*, 2017) | Secreted chicken netrin concentrated from media | 250ng/ml | 24 hours in 2 DIV cultures | Cortical neurons from mouse (E15) | TRIM9 negatively regulates FAK activity and netrin-dependent axon branching. |
| This study | Secreted mouse netrin concentrated from media | 600 ng/ml and 1200ng/ml | For 24 hours in 2DIV cultures | Cortical neurons from mouse (E15) | TRIM9 regulate concentration-dependent biphasic netrin response. FAK activity is increased in response to high netrin-1 concentration. |

**Table 1. Summary of the effect of netrin-1 on FAK activity in vitro.**

Li, W., Lee, J., Vikis, H.G., Lee, S.-H., Liu, G., Aurandt, J., Shen, T.-L., Fearon, E.R., Guan, J.-L., and Han, M. (2004). Activation of FAK and Src are receptor-proximal events required for netrin signaling. Nature neuroscience *7*, 1213-1221.

Liu, G., Beggs, H., Jürgensen, C., Park, H.-T., Tang, H., Gorski, J., Jones, K.R., Reichardt, L.F., Wu, J., and Rao, Y. (2004). Netrin requires focal adhesion kinase and Src family kinases for axon outgrowth and attraction. Nature neuroscience *7*, 1222-1232.

Moore, S.W., Zhang, X., Lynch, C.D., and Sheetz, M.P. (2012). Netrin-1 attracts axons through FAK-dependent mechanotransduction. Journal of Neuroscience *32*, 11574-11585.
Plooster, M., Menon, S., Winkle, C.C., Urbina, F.L., Monkiewicz, C., Phend, K.D., Weinberg, R.J., and Gupton, S.L. (2017). TRIM9-dependent ubiquitination of DCC constrains kinase signaling, exocytosis, and axon branching. Molecular biology of the cell *28*, 2374-2385.
Ren, X.-r., Ming, G.-l., Xie, Y., Hong, Y., Sun, D.-m., Zhao, Z.-q., Feng, Z., Wang, Q., Shim, S., and Chen, Z.-f. (2004). Focal adhesion kinase in netrin-1 signaling. Nature neuroscience *7*, 1204-1212.
Tang, F., and Kalil, K. (2005). Netrin-1 induces axon branching in developing cortical neurons by frequency-dependent calcium signaling pathways. Journal of Neuroscience *25*, 6702-6715.
